# Natural Selection on MHC IIb in Parapatric Lake and Stream Stickleback: Balancing, Divergent, Both, or Neither?

**DOI:** 10.1101/096917

**Authors:** William E. Stutz, Daniel I. Bolnick

## Abstract

Major histocompatibility (MHC) genes encode proteins that play a central role in vertebrates’ adaptive immunity to parasites. MHC loci are among the most polymorphic in vertebrates’ genomes, inspiring many studies to identify evolutionary processes driving MHC polymorphism within populations, and divergence between populations. Leading hypotheses include balancing selection favoring rare alleles within populations, and spatially divergent selection. These hypotheses do not always produce diagnosably distinct predictions, causing many studies of MHC to yield inconsistent or ambiguous results. We suggest a novel strategy to distinguish balancing versus divergent selection on MHC, taking advantage of natural admixture between parapatric populations. With divergent selection, immigrant alleles will be more infected and less fit because they are susceptible to novel parasites in their new habitat. With balancing selection, locally-rare immigrant alleles will be more fit (less infected). We tested these contrasting predictions using threespine stickleback from three replicate pairs of parapatric lake and stream habitats. We found numerous positive and negative associations between particular MHC IIβ alleles and particular parasite taxa. A few allele-parasite comparisons supported balancing selection, others supported divergent selection between habitats. But, there was no overall tendency for fish with immigrant MHC alleles to be more or less heavily infected. Instead, locally rare MHC alleles (not necessarily immigrants) were associated with heavier infections. Our results illustrate the complex relationship between MHC IIβ allelic variation and spatially varying multi-species parasite communities: different hypotheses may be concurrently true for different allele-parasite combinations.

## Introduction

MHC class II loci, which aid in the recognition of extracellular parasites, are among the most polymorphic loci in vertebrates’ genomes (Figueroa *et al.* 1988). Evolutionary biologists have long sought to elucidate the evolutionary processes that maintain the exceptional diversity of MHC within and among populations. Most studies have focused on documenting parasite-mediated selection on MHC given its role in immunity. Parasite-derived proteins (antigens) are collected and fragmented by antigen presenting cells (Roche & Furuta 2015). MHC proteins bind to certain antigen sequences, and export these to the cell surface for presentation to T-cells, which may then initiate an immune response. MHC II β chains with different peptide binding region sequences enable recognition of different parasite antigens (Eizaguirre & Lenz 2010; Hedrick 2002). Accordingly, MHC polymorphism contributes to variation in resistance to parasites including pathogenic and symbiotic bacteria (Bolnick *et al.* 2014; Kubinak *et al.* 2015; Lohm *et al.* 2002), viruses (Thursz *et al.* 1995), protozoa (Hill *et al.* 1991; Sinigaglia *et al.* 1988; Wedekind *et al.* 2006), helminthes (Paterson *et al.* 1998), fungi (Savage & Zamudio 2011), and even contagious cancers (Siddle *et al.* 2010). Despite these and many other studies, it remains unclear how MHC polymorphism is sustained. The leading hypotheses invoke balancing selection within populations, or divergent selection among populations, each of which has received mixed support (Bernatchez & Landry 2003; Piertney & Oliver 2006; Tobler *et al.* 2014; Yasukochi & Satta 2013).

Balancing selection occurs when rare alleles gain an inherent fitness advantage over common alleles, preventing their loss and maintaining allelic diversity (Takahata & Nei 1990; Takahata *et al.* 1992). Balancing selection can result from heterozygote advantage because individuals carrying more diverse MHC alleles recognize and resist more diverse parasites (Doherty & Zinkernagel 1975; Oliver *et al.* 2009), thanks to co-dominance (Lohm *et al.* 2002). Because rare alleles tend to occur in heterozygotes, they increase fitness and are protected from loss (Wegner *et al.* 2003). Alternatively, balancing selection can result from negative frequency-dependent selection. Parasites evolve strategies to exploit locally common host genotypes, such as evading detection by locally common MHC alleles (Slade & McCallum 1992). Because rare alleles do not provoke parasite counter-evolution, they may be more effective at detecting and protecting against local parasites (Muirhead 2001; Schierup *et al.* 2000).

Divergent natural selection (divergent selection) is also widely invoked to explain MHC diversity (Hedrick 2002; Hill *et al.* 1991; Meyer & Thomson 2001). Parasite communities often differ among host populations, favoring different MHC alleles in different locations and driving between-population divergence but undermining local polymorphism. Many studies have invoked divergent selection on MHC to explain allele frequency differences between populations with different parasites (e.g., (Copley *et al.* 2007; Matthews *et al.* 2010; Pavey *et al.* 2013). But, many studies do not formally test the null hypothesis that MHC divergence is neutral and unrelated to parasitism (Miller *et al.* 2010). Those that do consider neutrality often find mixed results: MHC divergence sometimes is greater than, less than, or equal to neutral genetic markers (Lamaze *et al.* 2014; Mona *et al.* 2008; Schwensow *et al.* 2007; Sutton *et al.* 2011). An alternative approach to test divergent selection is to evaluate whether different MHC alleles confer protection in different populations, using spatial variation in MHC-parasite associations to argue for divergent selection (e.g. (Eizaguirre *et al.* 2012a; Loiseau *et al.* 2009). Comparatively few studies have used experimental transplants or infections to test for of local adaptation at MHC loci (Eizaguirre *et al.* 2012a, b; Evans *et al.* 2010), and some of these have yielded negative results (Rauch *et al.* 2006).

Unfortunately, divergent and balancing selection may be difficult to distinguish because in certain contexts they can result in similar patterns, as pointed out by Spurgin and Richardson (2010) and more recently by Tobler et al. (2014). Both heterozygote advantage and negative frequency-dependent selection can lead to fluctuating allele frequencies through time (Slade & McCallum 1992). If these allele frequency fluctuations are asynchronous across host populations (Gandon 2002), then populations will be genetically divergent. Experimental transplants between such populations may (transiently) yield signals that appear to support divergent selection, even though MHC divergence arose from balancing selection within populations. Still more problematic, balancing and divergent selection are not mutually exclusive phenomena. Balancing selection may act within populations (driven by some parasites), while other parasites generate divergent selection favoring differences between populations. Simultaneous balancing and divergent selection may obscure each force’s effect on within-population diversity and between-population divergence. Lastly, the majority of studies using MHC-parasite associations to test for selection have focused on one parasite species at a time. This inevitably yields an incomplete picture of the selective forces shaping diversity at MHC, especially because different parasites may drive different kinds of selection. Consequently, tests for balancing selection and divergent selection have yielded mixed evidence (Eizaguirre & Lenz 2010; Spurgin & Richardson 2010; Yasukochi & Satta 2013).

### Balancing versus divergent selection in parapatry

In certain settings, balancing and divergent natural selection can lead to unique and thus testable outcomes. In particular, we suggest they can be distinguished in parapatric populations that actively exchange migrants but experience distinct parasite communities (Fig. 1), by estimating three parameters. δ_m_ measures how strongly an allele *m* is enriched in a focal habitat. θ_p_, measures how strongly a parasite taxon *p* is enriched in a focal habitat. β_mp_ measures the association between allele *m* and parasite *p;* negative values imply that the presence of the allele coincides with lower parasite abundance (Fig. 1A).

**Figure 1.**
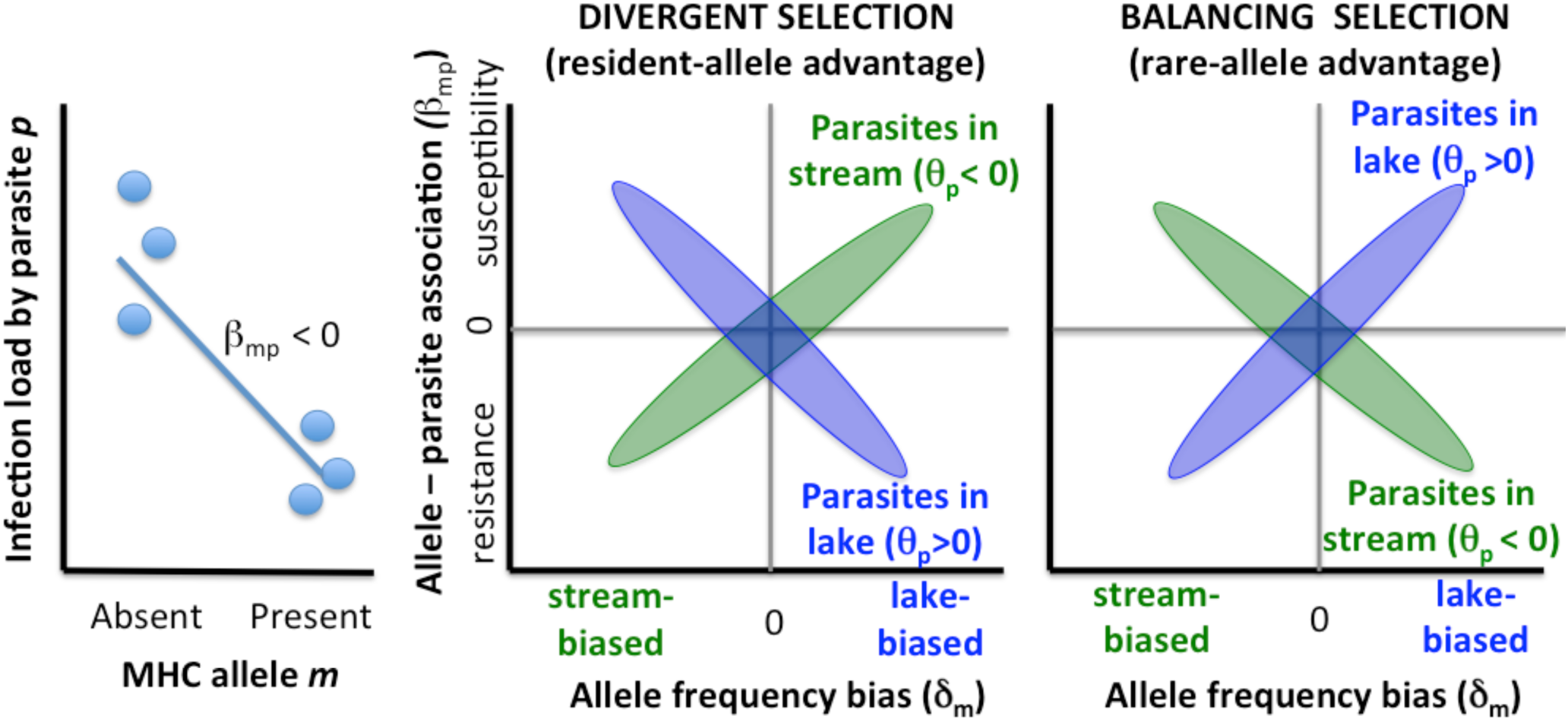
A conceptual diagram illustrating our strategy to test for a predominant effect of divergent selection between populations, versus balancing selection within populations. We calculate an effect size and direction (β_mp_) for all pairwise associations between MHC allele *m* versus parasite taxon p, within a given habitat (lake or stream). Then, we test whether β_mp_ depends on the extent to which allele *m* is lake- or stream-biased δ_m_), and the parasite is lake- or stream-biased (θ_p_). We expect that β_mp_ covaries with δ_m_, but the sign of this trend is opposite within the lake versus stream samples. With divergent selection, each population will contain locally common alleles that confer protection against locally common parasites, whereas immigrants will tend to be susceptible to unfamiliar parasites. For example, alleles that are particularly common in the lake (δ_m_>0) should confer protection (β_mp_<0) against lake-specific parasites (θ_p_>0), but susceptibility (β_mp_>0) against parasites in the stream (θ_p_<0). In contrast, balancing selection favors rare alleles, so immigrant alleles should benefit. Stream-biased MHC alleles (δ_m_<0) that migrate into the neighboring lake should be rare and confer resistance (β_mp_<0) to lake-specific parasites (θ_p_>0). Therefore, both divergent and balancing selection should produce a θ_p_*δ_m_ interaction effect on β_mp_, but the direction of this interaction depends on the form of selection.

Divergent selection will tend to increase the abundance of an allele in the habitat where it confers a protective benefit (β_mp<_0), or decrease the allele in a habitat where it confers susceptibility (β_mp>_0). Consequently, alleles that are strongly enriched in a particular habitat (δ_i_) should tend to be protective (β_mp<_0) against parasites enriched in that same habitat (θ_p_,>0). In the context of our study system (lake and stream populations of threespine stickleback, details below), this means that the more lake-biased alleles should protect against lake-biased parasites (and be susceptible to stream-biased parasites). Conversely, stream-biased alleles should protect against stream-biased parasites and be susceptible to typical lake parasites (Fig. 1B).

Balancing selection will tend to favor alleles that are locally rare, which in parapatric settings includes immigrants. Namely, when there is balancing selection we expect that alleles enriched in a particular habitat (relative to the neighboring habitat) will be particularly susceptible to parasites from that habitat (β_mp>_>0; Fig. 1C). In contrast, alleles that are scarce in a focal habitat will tend to protect against the local parasites (Muirhead 2001; Schierup *et al.* 2000). These locally rare alleles could be new mutations or (more frequently in a parapatric setting) immigrants (Lamaze *et al.* 2014). In this regard, balancing selection resembles local maladaptation, the diametric opposite of expectations for divergent selection.

Thus, divergent and balancing selection make opposite predictions regarding the sign of the correlation between δ_m_ (the extent to which an allele is habitat-specific) and β_mp_ (the allele’s effect on parasites), for habitat-biased parasites (θ_p_ Fig. 1). Do endemic macroparasites disproportionately infect hosts with locally-enriched alleles (implying balancing selection) or locally-depleted alleles (implying divergent selection)? To test these predictions we rely on natural migrants between populations, which add rare genetic variants that are either beneficial (balancing selection), or deleterious (divergent selection). Of course, both selective forces might act concurrently, for instance if certain parasites select against immigrants while other parasites select against locally common alleles. Therefore, any test of these alternative predictions should take into account the full set of MHC alleles, and all common parasites within each population.

Some previous studies have used a related approach, testing whether MHC alleles confer protection or susceptibility to different parasites in different habitats (e.g., estimating β_mp_) (Tobler *et al.* 2014). But, a key element of our approach is that the sign and strength of MHC-parasite associations (β_mp_) will depend on the extent of between-population differences in parasite and allele frequencies (θ_p_, and δ_m_). To our knowledge, previous studies of MHC adaptation have not tested for an interactive effect of θ_p_, and δ_m_ on β_mp_ (parasite-habitat and allele-habitat biases jointly affecting the parasite-allele association). Here, we use this novel approach to test for signatures of balancing or divergent selection in connected (parapatric) lake and stream populations of threespine stickleback (*Gasterosteus aculeatus*).

### Study system: threespine stickleback

Genetic sequencing suggests that threespine stickleback have between 4 and 6 functional MHC class IIβ loci in their genome (Reusch *et al.* 2004; Reusch & Langefors 2005; Sato *et al.* 1998), though this may vary between individuals (Reusch & Langefors 2005). Expression analysis indicates that all putative MHC class IIβ loci are typically expressed (Reusch *et al.* 2004).

Prior studies of stickleback have provided evidence for balancing or divergent selection on MHC IIβ. Balancing selection is supported by several observations. Individuals with an intermediate number of alleles are more resistant to infection (Kurtz *et al.* 2004; Wegner *et al.* 2004), harbor fewer parasites (Wegner *et al.* 2003), build better –quality nests (Jager *et al.* 2007), survive better (McCairns *et al.* 2011; Wegner *et al.* 2008), and attain higher lifetime reproductive success (Kalbe *et al.* 2009). Divergent selection is supported because MHC IIβ allele differ between (i) co-occurring benthic and limnetic stickleback species pairs (Matthews *et al.* 2010), (ii) closely parapatric estuarine stickleback in Quebec (McCairns *et al.* 2011), and (iii) lake and river stickleback from northern Germany (Rauch *et al.* 2006; Reusch *et al.* 2001). The German lake/river system has been used for experimental tests of divergent selection on MHC IIβ. Lab-bred F2 lake-stream hybrids placed into field mesocosms gained more weight if they had local MHC alleles, but non-native MHC genotypes were not systematically more infected (Eizaguirre *et al.* 2012a). An earlier F2 hybrid transplant experiment found that genomic background but not MHC genotype explained habitat-specific infection rates (Rauch *et al.* 2006).

Here, we present a simultaneous test both balancing and divergent natural selection on stickleback MHC IIβ in three replicate lake-stream pairs of stickleback. We first document between-habitat differences in parasite composition (θ_j_), and MHC genotypes (δ_i_). Then, for each of the three pairs, we test for associations between each MHC IIβ and each parasite taxon (β_ij_) within a multispecies parasite community. Lastly, we test whether allele-parasites associations covary positively or negatively with habitat differences in allele and parasite frequencies (Fig. 1). Specifically, we test whether locally common parasites disproportionately infect locally-enriched alleles, or locally-rare immigrant alleles.

## Methods

### Collections

In July 2007, we sampled threespine stickleback from three lakes on northern Vancouver island, British Columbia (Roberts Lake, Farewell Lake, and Comida Lake) and their corresponding outlet streams (three ‘lake-stream pairs’, Fig. 2). Most lake-stream pairs on Vancouver Island evolved independently *in situ*, after marine stickleback colonized freshwater after Pleistocene deglaciation (Clague & James 2002; Hendry *et al.* 2013; Stuart *et al.* In review).

We collected adult stickleback using unbaited minnow traps (0.5-cm gauge). We placed traps haphazardly along the shoreline of each lake (< 3m depth) within 350 meters of the outlet stream, and at 5 traps at each of multiple locations along each lake’s outlet stream (Table 1, Fig. 2). Stream samples spanned the genetic clinal transition from lake- to stream-genotypes (Berner *et al.* 2009; Weber *et al.* 2017). Upon capture, fish were immediately euthanized in MS-222. Caudal fin clips were taken and preserved in 90% ethanol for later DNA extraction. Fish were preserved in 10% neutral buffered formalin. Collection and animal handling were approved by the University of Texas Institutional Animal Use and Care Committee (Protocol # 07-032201), and a Scientific Fish Collection Permit from the Ministry of the Environment of British Columbia (NA07-32612).

**Figure 2.**
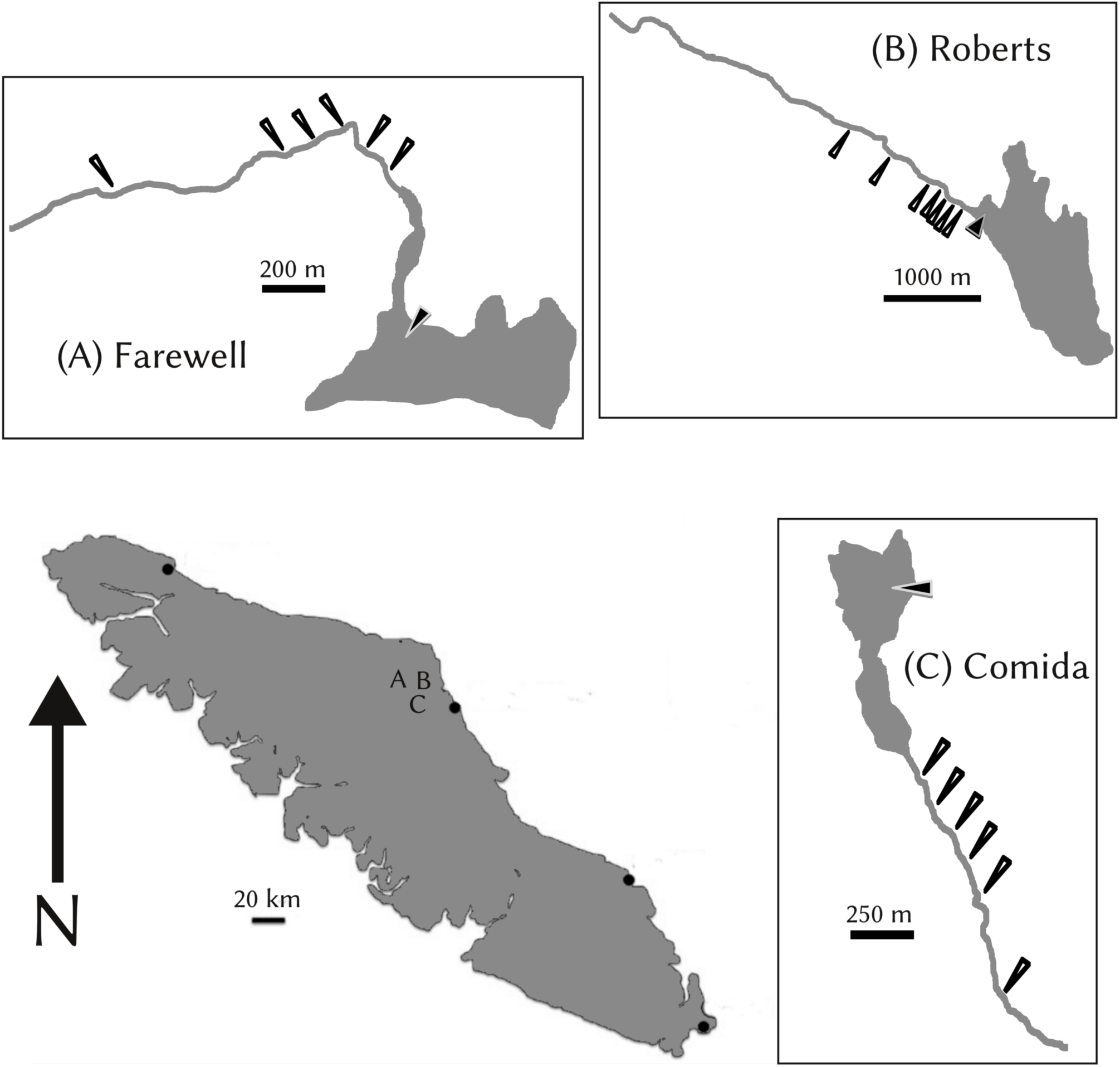
Map of study system showing Vancouver Island and the three lake-stream pairs used in this study. Approximate locations of each lake-stream pair on the island are indicated by their respective letters (A: Farewell, B: Roberts, C: Comida). Arrows indicate separate sampling locations within each pair. Separate scale bars are provided for the entire island and each pair individually.

### Parasite load

Each fish was exhaustively screened to enumerate macro-parasites (helminths, crustaceans, molluscs, and microsporida) visible under a standard dissection microscope. This included scans of the outer body (i.e. skin and bony armour structures), mouth and gills, interior body cavity including all organs (liver, swim bladder, gonads), the interior of the intestinal tract (stomach and intestine), and the eyes (interior and exterior). Only the gills on the right (but not left) side of the fish were scanned for parasites, as the common gill parasites (*Thersitina* sp. and *Unionidae* glochidia) were present at very high abundances on both left and right gills. All parasites were identified to the lowest possible taxonomic unit (genus in most cases).

### Analysis of habitat effect on infection

To determine whether parasite abundance differed between lake and stream habitats, we first fit hierarchical generalized linear models separately for each lake stream pair. The GLMs used (additively) overdispersed-Poisson distributions to model each parasite taxon’s abundance in individual fish. The basic form of each model was:

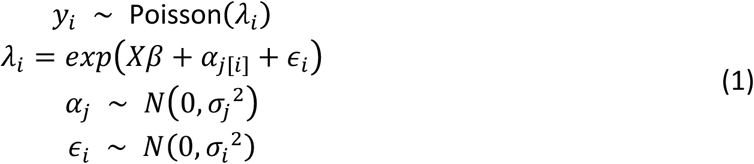

where y_i_ is the abundance of a focal parasite taxon in individual *i*. The term α_j_ denotes a habitat-specific intercept where *j*=lake, or stream. The vector β includes the regression parameters β_1_ through β_5_ which indicate, respectively, the means for lake (β_1_) and stream (β_2_) habitat, the covariate effect of fish standard length of lake fish (β_3_) and stream fish (β_4_), and a coefficient for sex (β_5_). Random effects (α_i_) associated with each sampled stream site (i.e. 100m, 200m, etc.), are modeled as a normal random variable with mean equal to zero and standard deviation σ*_j_*. The error terms ε_i_ that account for overdispersion in the abundance data were also modeled as normal random variables with mean equal to zero and standard deviation σ*_i_*. Sex was centered at zero, and length was centered at zero prior to fitting the models (Gelman & Hill 2006). Thus, the models explicitly account for the effects of sex, size, and heterogeneity among sampling locations (e.g., within-stream clines) and sample sizes when estimating mean abundances within each habitat (β_1_ and β_2_).

All parameters were estimated by drawing 1000 samples from their joint posterior distributions using the Markov Chain Monte Carlo (MCMC) algorithm implemented the *MCMCglmm* package (Hadfield 2010) in R version 3.2.1. Weakly informative normal priors with a scale of 3 and 10 were applied to all fixed slope and intercept coefficients respectively, providing some shrinkage of β estimates away from extremely large values (Gelman *et al.* 2008). Half-Cauchy priors with scale equal to 10 were applied to α_j_’s, while a uniform prior was applied to the residual standard deviation α_i_. In cases where hyperparameter variances were close to zero, stronger half-Cauchy or inverse-Wishart priors were used to improve model convergence. MCMC chain parameters were determined heuristically by increasing the thinning interval until all estimated parameters achieved an autocorrelation less than 0.1.

As our metric of parasite habitat bias we calculated the posterior distributions for a derived parameter (θ_p_), which was the log of the ratio of parasite *p*’*s* mean abundance estimates (on the data scale) between the lake and the stream. When θ_p_>0, the focal parasite is more abundant in the lake, and when θ_p_<0 the parasite is more abundant in the stream. Parasites with greater than 95% percent posterior probability of being at least two times more abundant in one habitat that were considered strongly 'habitat-specific’ in subsequent analyses. We use ‘habitat-biased’ to refer to weaker habitat effects.

### MHC sequencing and genotyping

We genotyped MHC IIβ from a random subset of the fish that were screened for parasites (sample sizes listed in Table 2), by 454 pyrosequencing of PCR amplicons. The procedures for DNA extraction, quantitation, PCR amplification, and library preparation, and computational analysis are described fully in (Stutz & Bolnick 2014). We used PCR primers that produce a 210 base pair amplicon (excluding primer sequences) covering 75% of the length of exon 2 (210 bp out of 265 bp of exon2; (Stutz & Bolnick 2014). This covers 70 out of 88 amino acid residues, including the highly variable peptide binding region (PBR) of the exon (Lenz et al. 2009a). Of the 846 fish genotyped in the present study, 295 were previously described in Stutz and Bolnick (2014). We genotyped the additional samples in four new pyrosequencing runs (1/4 plate per run).

Our analytical pipeline uses a quasi-Dirichlet process to iteratively cluster similar sequence reads into groups at increasing levels of sequence similarity, and estimates whether clusters represent single true allelic variants present in the original sample (Stutz & Bolnick 2014). A separate research group independently tested this bioinformatics pipeline, using multiple datasets, and confirmed its accuracy (Sebastian *et al.* 2016). Allelic sequences for each individual were aligned to the cloned sequences in Sato et al. (1998) to ascertain phase, then translated into amino acid sequences for further analysis. Hereafter we refer to a unique amino acid sequence as an ‘allele’. We focus on allele presence or absence, because MHC is expected to have co-dominant effects on parasites (Doherty & Zinkernagel 1975).

### Analysis of habitat effect on MHC genotype

We applied a similar hierarchical modeling approach estimate allele frequency bias between habitats within each lake-stream pair. Because an MHC allele may be distributed across multiple paralogs, this is not a traditional allele frequency, but rather the proportion of fish carrying an allele. For each allele we fit the following model:

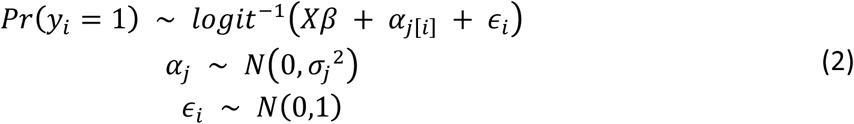

where *y_i_*=1 indicates that fish *i* carries the allele. The vector β contains separate intercept coefficients for the lake and stream (β_1_ and β_2_) as well as coefficients for sex (β_3_) and size (β_4_, β_5_) while the *α_j_* term indicates additional (random) effects associated with each sampled stream site *j*. The variance of ε_i_ was fixed at one due to non-identifiability of individual-level overdispersion in binomial GLMs (Gelman & Hill 2006). As with parasites, non- or weakly informative priors were used for all parameters, which were estimated by drawing 1000 samples from their joint posterior distributions using *MCMCglmm*.

As a metric of habitat bias (whether allele frequency was greater in one habitat or the other), we estimated the derived parameter δ*_m_* for each allele *m*, which is equal to the log of the ratio of allele frequency estimates for the lake and the stream. Alleles with δ<0 are more common in the lake, and δ<0 are more common in the stream. Alleles with a 95% posterior probability of occurring at least twice as frequently in one habitat were considered ‘habitat-specific’ in subsequent analyses. We use 'habitat-bias' to refer to a less stringent form of divergence (e.g., 95% posterior for δ excludes 0).

### Estimating MHC allele effects on infection

We next estimated whether the presence or absence of each MHC allele *m* in individual fish is associated with each parasite taxon *p*’*s* abundance. If an allele helps the host recognize and resist a particular parasite, then individuals with that allele should be less intensely infected by that parasite. We call this a 'negative' allele-parasite association because the allele has a negative effect on the parasite. Positive associations (an allele’s presence coincides with heavier infection loads) can arise for several reasons including susceptibility, if the parasite directly exploits that allele to establish an infection (Westerdahl *et al.* 2012).

For each lake-stream pair, we used hierarchical models to estimate the effect of each MHC allele on each parasite. The basic form of each model was:

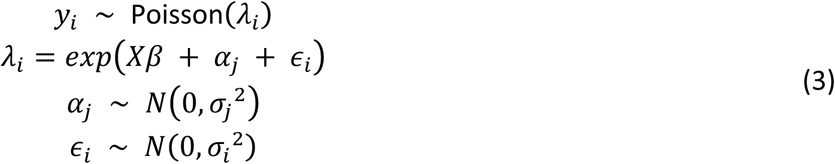

where y_i_ is the abundance of a given parasite taxon *p* in individual *i.* The vector β includes the same 5 regression coefficients as the parasite specificity models, plus an additional coefficient associated with the presence/absence of the focal allele *m* (β_6_). As before, the *α_j_* term indicates samplinglocation effects on abundance (i.e. 'random' effects). The error term ε_i_ gives the fitted residual error for individual *i*, accounting for any observed overdispersion in the abundance data. For the few instances where one MHC allele strongly covaried with another allele (Yule’s |Q| > 0.8, see Supplementary Material), we included the correlated (non-focal) allele as an additional factor in our model. When two alleles were perfectly correlated, however, we dropped the less common allele.

Models were fit in a Bayesian probability framework using MCMC sampling implemented in the *MCMCglmm* package (Hadfield 2010). Alleles were transformed from 0/1 variables to a continuous variable with a mean zero to avoid issues with separation. Fish length was scaled to a mean of zero and standard deviation of 1 (within habitats) prior to model fitting (Gelman *et al.* 2008). As before, MCMC chain parameters were determined heuristically by increasing the thinning interval until all estimated parameters achieved an autocorrelation less than 0.1. Posteriors for each allele/parasite combination were estimated by drawing 1000 samples from their joint posterior distributions. Posterior means, standard errors, and 95% high probability density intervals (HPDIs) for all estimated allele effect sizes were calculated from these posterior distributions.

We use the parameter β_6_ as our measure of the effect of allele *m* on parasite *p*, which we hereafter denote β_m_,_p_. Because we assume that most true allele effects are zero (a given allele has no discernible effect on a given parasite), but that a few alleles will have moderate to strong effects on parasite abundance, those effects whose 95% HPDI's for β_m_,_p_ did not include zero were delineated as "non-zero" effects. Note that when β_m_,_p_>0, the focal allele is associated with higher abundance of the given parasite in a given habitat, and when β_m_,_p_<0 the allele is associated with lower abundance. For shorthand we refer to these alternative outcomes as susceptibility and resistance, respectively.

### Testing for balancing or divergent selection

We used the estimates of MHC allele frequency differences between habitats (δ_m_), parasite abundance differences between habitats (θ_p_) and MHC-parasite association strengths (β_mp_ coefficient in model (3) above), to test for signatures of balancing or divergent selection using the logic explained in the introduction (Fig. 1). Specifically, we tested whether lake-biased alleles (δ_m_>0) disproportionately protect the host (β*_mp_*<0) from lake-biased parasites (θ_p_> 0), and conversely whether stream-biased alleles (δ_m_<0) protect the host (β*_mp_*<0) from stream-specific parasites (θ_p_<0). When defining stream-specific parasites (or alleles), we retain those whose 95% HPDI of θ_p_ (or δ_m_) excludes zero. Our prediction can be tested with a linear model examining whether alleles’ protective effects (β*_mp_*) depend on an interaction between the allele-frequency bias (δ_m_) and which habitat a parasite is specific to (θ_p_). Balancing selection should also generate a δ_m_*θ_p_ interaction, but with the opposite slopes compared to divergent selection (Fig. 1C).

The above analysis focuses only on strongly lake- and stream-specific parasites, because these are most likely to drive divergent selection. As a consequence, that analysis omits parasites that are common or rare in both habitats. We repeated the analysis by regressing each MHC-parasite effect (β*_mp_*) on the relevant allele’s habitat bias (δ_m_), parasite’s habitat bias (θ_p_), and a δ_m_*θ_p_ interaction, with habitat as a factor as well. We expected to observe a significant δ_m_*θ_p_ interaction whose direction would distinguish between selection models.

The preceding tests focus on allele frequency differences between habitats (δ_m_), rather than absolute allele frequencies within habitats. This is most appropriate when considering gene flow and divergent selection, but local absolute allele frequency may be more relevant to frequency-dependent selection by parasites. We therefore repeated the analyses described above, but using within-habitat MHC allele frequency instead of the between-habitat frequency difference (δ_m_). Specifically, we regressed the MHC-parasite effect β*_mp_* against the allele’s frequency in whichever habitat the focal parasite is most abundant in (e.g., stream frequency when θ_p_<0, lake frequency when θ_p_>0). We did this focusing on only the convincingly non-zero MHC-parasite associations (whose 95% HDPI excludes zero), and then again using all pairwise associations.

## Results

### Parasite abundance differences between habitats

A total of 34 parasite taxa were identified across the three lake-stream pairs, although not every parasite was present in every population or pair (Fig. 3). Within each pair, the parasite community differed substantially between habitats. Per-fish parasite richness was significantly higher in lake than stream habitats for all three pairs (Fig. S1). More parasite taxa were strongly habitat-specific to lakes (θ_p_ >> 0, for 8, 5, and 5 taxa in Comida, Farewell, Roberts Lakes respectively) than to their adjoining streams (2, 0, and 0 taxa respectively). *Crepidostomum* was the only parasite that was considered lake-specific in all three lake-stream pairs (Blackspot, *Thersitina*, and *Unionidae* were lake-specific in two of three pairs). *Anisakis* and *Bunodera* met our approaches our strict habitat-specific threshold in the three streams. But, most parasites exhibit variable, weak, or no habitat affiliation (Fig. 3).

**Figure 3.**
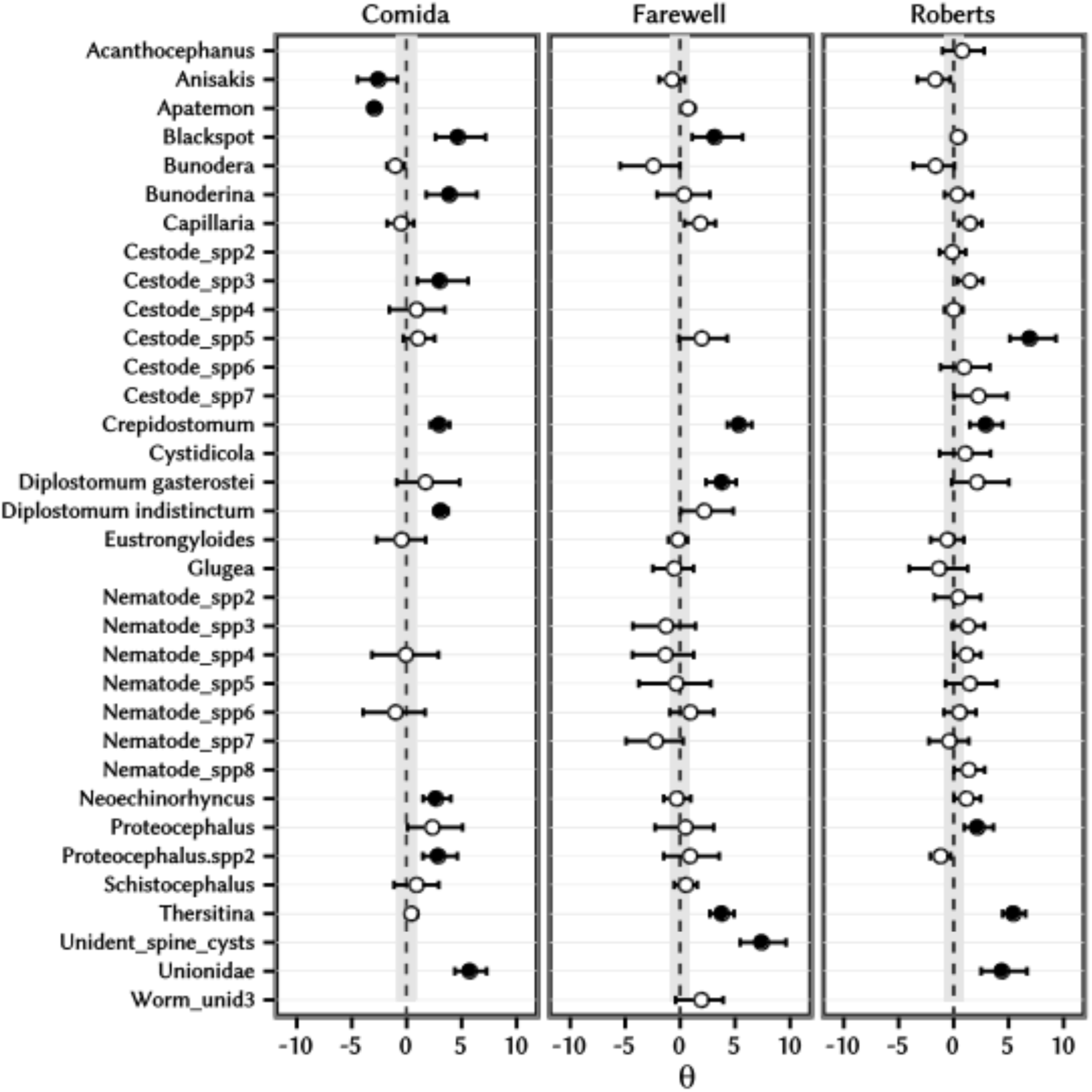
Posterior distributions of parasite prevalence differences between habitats (θ_p_) for each parasite within each for lake-stream pair indicate which parasites are habitat specific. Positive values of θ_p_ indicate lake-biased parasites, while negative values indicate stream bias. Note θ_p_ is calculated on a logarithmic scale. Posterior means are indicated by circles while the bars indicate 95% credible intervals. Credible intervals must fall completely above or below the gray band in each panel to meet our criterion of regarding parasites as habitat-specific (a high probability of being at least twice as abundant in one habitat than in the other). Filled circles indicate habitat-specific parasites.

### Allele prevalence differences between habitats

We identified 374 unique MHC alleles across our three lake-stream pairs, 95% of which were restricted to a single lake-stream pair. Within each lake-stream pair, up to 13% of the MHC alleles were strongly habitat-specific specific (at least a 2-fold frequency difference, |δ_m_|>>0; Fig. 4). There were more lake-specific alleles (9,7, and 9 in Comida, Farewell, and Roberts respectively) than stream-specific (2, 6, 2 respectively). No allele was habitat-specific in more than one lake-stream pair. For the 19 alleles shared among replicate pairs we found no parallel evolution of habitat differences (e.g., δ_m_ was not correlated across independent pairs).

**Figure 4.**
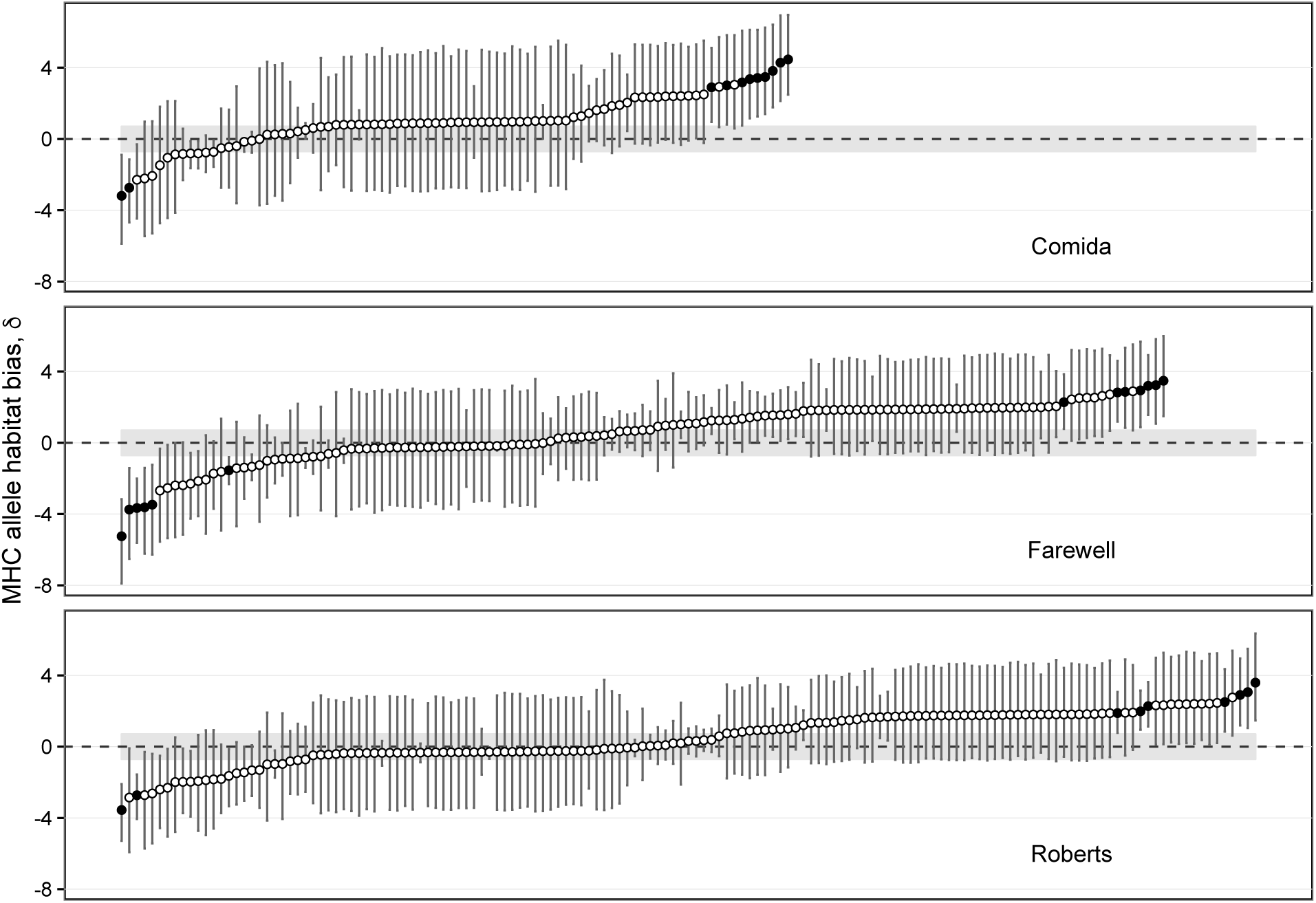
Posterior distributions of MHC IIβ allele differences between habitats (δ_m_) within each lake-stream pair. Positive values of δ_m_ indicate lake-biased alleles, while negative values indicate stream-biased alleles. Note that δ_m_ is calculated on a logarithmic (non-linear) scale. Posterior means are indicated by circles while the bars indicate 90% credible intervals. Credible intervals must fall complete above or below the gray band in each panel to indicate alleles with high probability of occurring twice as frequently in one habitat compared to the other. Habitat-specific alleles are indicated by filled circles and are labeled. Alleles are ordered along the x axis by increasing values of δ_m_. Note that few alleles are shared between the three lake-stream pairs.

The diversity of MHC is comparable between habitats (Fig S2). All sites exhibit on average about six unique MHC amino acid sequences per fish, albeit with substantial among-individual variation. In contrast, neutral genomic SNPs exhibited consistently lower nucleotide diversity in stream than lake fish (Fig. S3). Consequently, relative to neutral expectations MHC diversity is relatively higher in stream stickleback than in lake stickleback.

### Associations between MHC alleles and parasite infection

We estimated association strengths (β*_mp_*) between a total of 6006 combinations of MHC allele versus parasite taxon. Models for an additional 678 possible allele-parasite combinations failed to converge adequately, usually due to low extreme rarity of the parasite or allele. For the few cases where two alleles are statistically strongly linked (see Supp. Mat. on Yule’s Q), we dropped one of the redundant alleles. These tests found a substantial number of robust associations. Sixty-two MHC allele – parasite taxon associations had 95% HPDIs that did not include zero (Figs. 4&5, Table 4). No single allele is strongly associated with more than one parasite.

Overall, there were approximately 50% more negative than positive effects estimated within each pair (Table 3). Negative associations imply that fish carrying the focal allele are less-heavily infected by the focal parasite. The bias towards negative effects may be an artifact of comparing rare alleles to rare parasites, so henceforth we restrict our attention to strongly supported associations. Of those strong effects, 23 were negative. For example, stream-biased allele P293 coincided with an 18 fold reduction (β=-2.87 [-5.17,-0.78]) in the abundance of the lake-specific parasite *Crepidostomum* in Comida Lake (Fig. S4). Another 39 strong effects were positive, for instance lake-specific allele P342, associated with an 16-fold increase (β=2.75, [1.25,4.11]) in (lake-specific) Blackspot infection loads.

Several of the strong MHC-parasite associations exhibit a pattern consistent with local adaptation via divergent selection (Table S4): rare alleles conferring susceptibility to local parasites.Fish carrying MHC allele P273 carry ~3-fold more *Unionidae* (Fig. S5), which may explain why this allele is less common in the lake (δ=-2.73 [-4.68,-1.15]) where *Unionidae* are 314-fold more abundant (θ=5.75, [4.38,7.27]). Conversely, two MHC alleles are rarer in Comida stream and confer susceptibility to stream-specific *Apatemon*.

Other strong MHC-parasite associations support balancing selection: common alleles conferring susceptibility to local parasites. MHC allele P231 is 12-fold more common in Roberts Lake than stream (δ=2.50 [0.84, 4.36]). In the lake, it confers a 4-fold greater probability of infection by a lake-specific cestode (Fig. S6; θ=6.93, [5.12, 9.31]). Similarly, allele P403 is 72-fold more common in the Comida Lake (δ=4.28 [2.13, 6.94]) where it confers 3-fold higher risk of infection by lake-specific *Crepidostomum* (θ=2.99, [2.14, 3.91]; Fig. S7). Assuming these parasites reduce host fitness, these associations favor locally rare alleles over the currently-abundant P231 or P403 alleles.

### Tests for divergent or balancing selection

Aggregating across many such allele-parasite associations, we found no overall trends towards towards divergent or balancing selection. Regressions revealed no significant effect of habitat-biased MHC allele frequency (δ_m_) on the posterior mean estimate of the strong MHC-parasite associations (β*_mp_*; Fig. 6A). There was no significant relationship for either lake-specific parasites (θ_p_>>0, t_40_=1.22, P=0.246), or stream-specific parasites (θ_p_<<0, t_18_=0.465, P=0.647). Lake-stream pair had no effect in either regression (P>0.2). Putting these regressions into a single ANCOVA, we confirmed that the slopes of β*_mp_* ~ δ_m_ were indistinguishable from zero for both lake- and stream-specific parasites (overall δ_m_ effect P=0.1876). There was no significant interaction between δ_m_ and parasite-habitat (P=0.733), thus refuting both hypotheses’ expectation of opposing slopes for lake- versus stream-specific parasites. The effect of δ_m_ and the δ_m_* habitat interaction were also non-significant if we expanded the analysis to include all MHC-parasite associations (β*_mp_*, regardless of their strength, using only habitat-specific parasites. We expanded this still further to use all 6006 β*_mp_* estimates (regardless of strength of δ_m_, θ_p_, or β*_mp_*) we still found no significant θ_m_*δ_p_ interaction (t_6680_=1.49, P=0.1365). The only significant effect was that all MHC alleles (regardless of δ_m_) were more susceptible to lake-biased parasites than stream-biased parasites (Fig. S8; positive effect of θ_p_ on β*_mp_*; t_6680_=2.47, P=0.0137).

**Figure 5.**
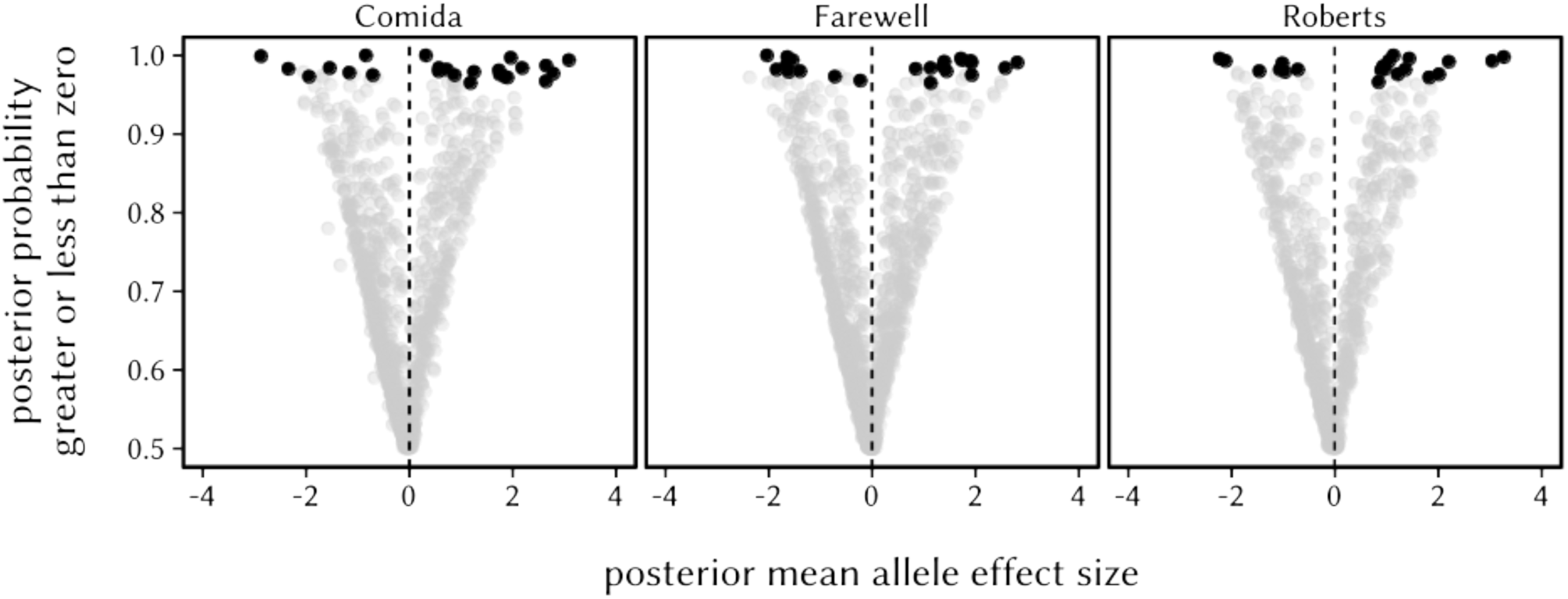
Volcano plot comparing the posterior mean effect sizes versus posterior probability of all MHC allele-parasite associations (β_mp_). Each point represents a single estimated effect of β_mp_, calculated for a given lake-stream pair. The y-axis shows the proportion of the posterior distribution greater than zero (for positive effects) or less than zero (for negative effects). Effect sizes are given on the latent (i.e. natural log) data scale. Black circles indicated effects with 95% HPD intervals that do not include zero.

**Figure 6.**
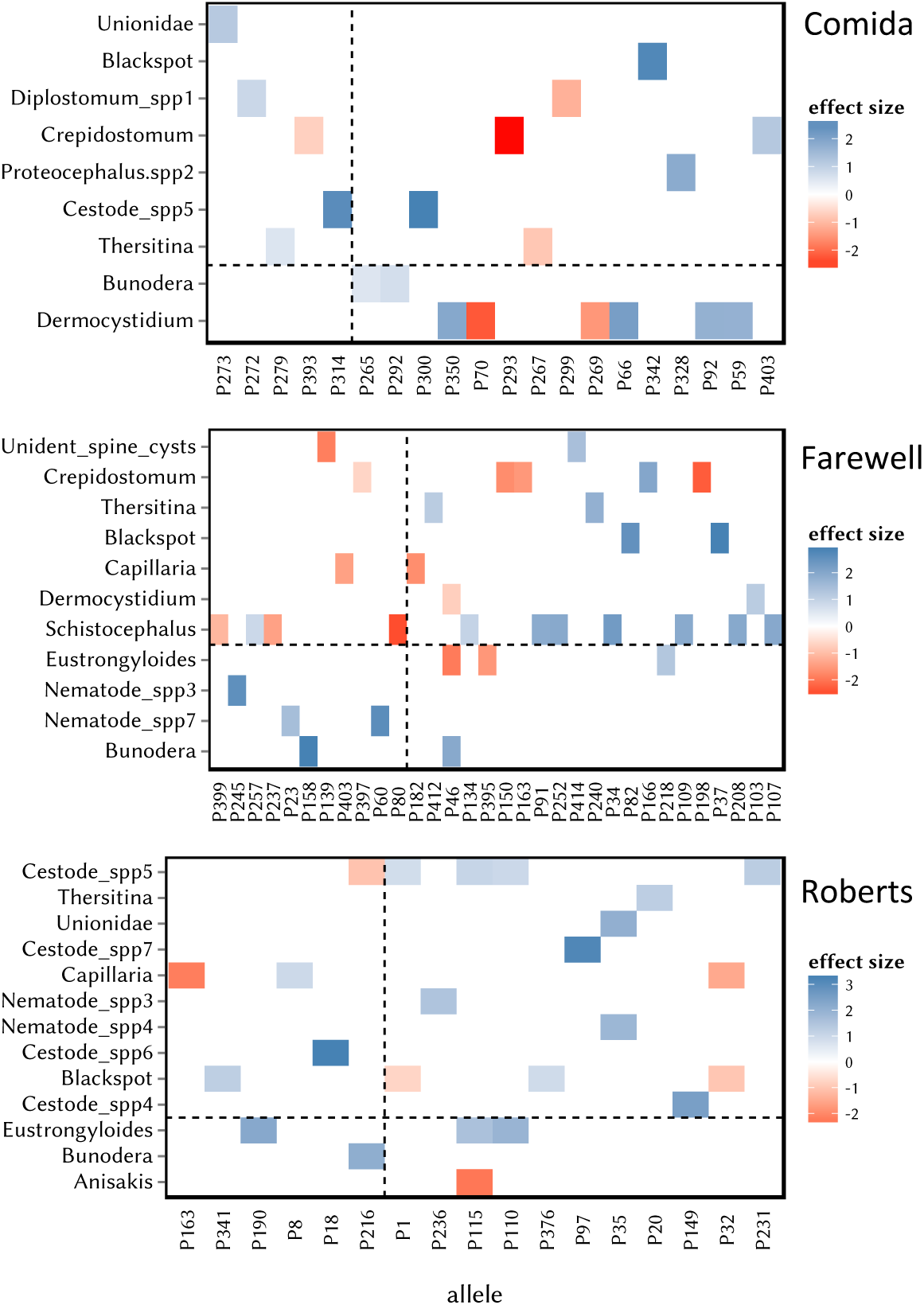
Heatmap of associations (β_mp_) between MHC alleles (x-axis) and parasite taxa (y-axis). We only plot associations whose 95% HPD intervals do not include zero (black circles from Fig. 5). Blue squares represent positive effects (alleles associated with higher parasite load), red represent negative effects (alleles conferring lower parasite load). We plot each lake-stream pair separately (Comida at the top, then Farewell, then Roberts). Within each pair, alleles to the right of the vertical dashed line are more common in the lake (δ_m_>0), alleles to the left are more common in the stream (δ_m_<0). Parasites above the horizontal dashed line are more common in the lake (θ_p_>0), parasites below the line are more common in the stream (θ_p_ <0). Examples of allele-parasite associations are plotted in the Supplementary Figures.

Lastly, we tested for effects of local allele frequency, rather than allele habitat-bias. MHC-parasite effect sizes (β*_mp_*) were negatively correlated with the focal allele’s frequency in the habitat where the parasite is relatively common (Fig. 7). This is true whether we focus only on large-effect estimates of β*_mp_* (t_60_=-3.87, P=0.00028), or on all estimates of β*_mp_* (t_6682_=-2.10, P=0.0446). For either variant on the analysis, the effects are weak (r^2^=0.186 and 0.0005, respectively). The negative trend arises because locally rare alleles (which are not necessarily immigrants) are most likely to confer susceptibility to local parasites (β*_mp_*>0). In contrast, locally common alleles are equally likely to exhibit positive or negative effects on local parasites (Fig. 7).

**Figure 7.**
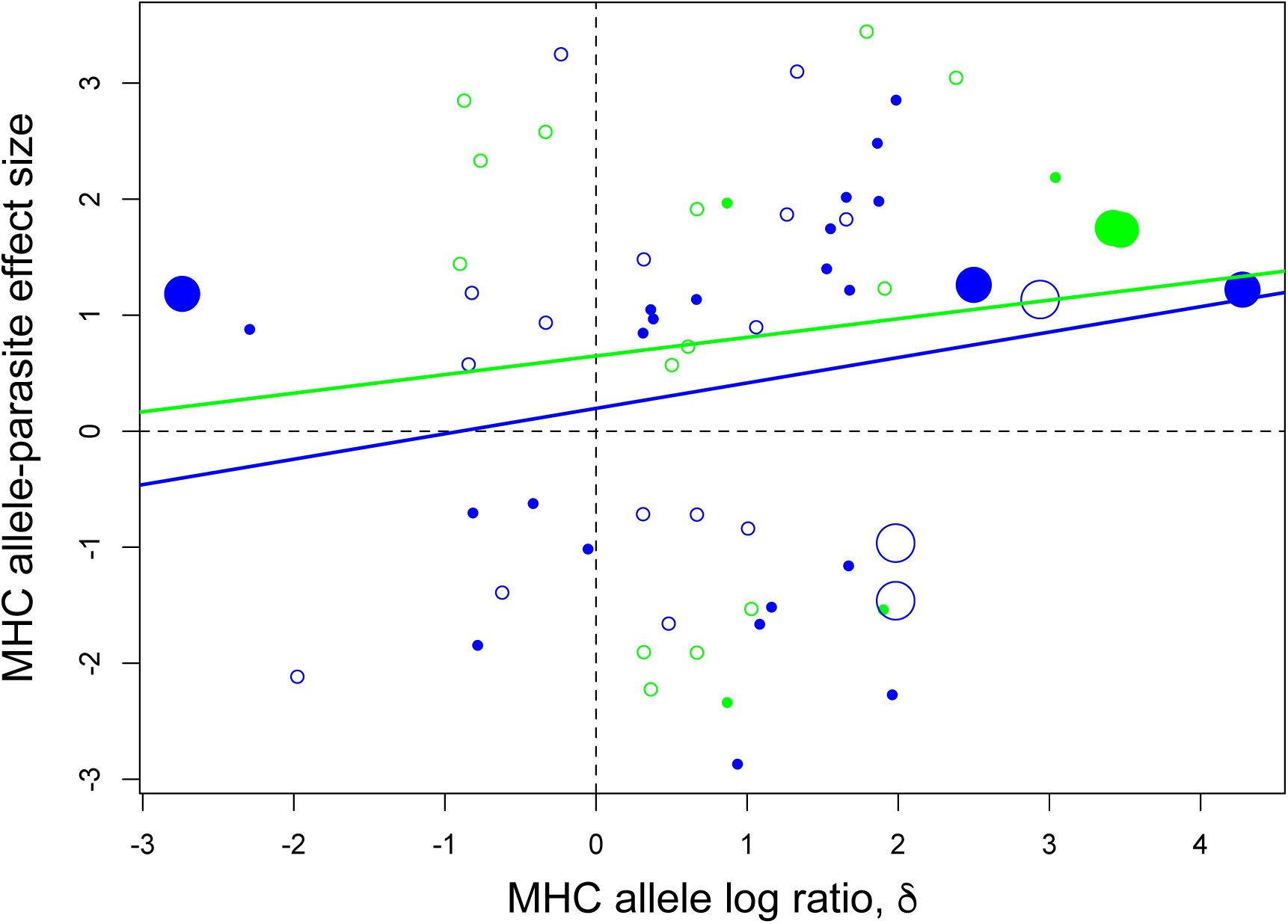
An empirical test of local adaptation versus balancing selection, as illustrated in our conceptual diagram (Fig. 1). We plot MHC allele effect size on parasites (β_mp_; positive values imply susceptibility, negative values imply resistance) as a function of the alleles’ relative abundance in the lake or stream (δ_m_; positive values imply higher frequency in the lake, and negative values imply higher frequency in the stream). Each point represents a non-zero association between an MHC allele and a parasite taxon (95% HPD intervals of β_mp_ do not include zero; black circles from Fig. 4). We plot separate regression lines for parasites that tend to be more common in the lake (blue, θ_p_>0) versus stream (green; θ_p_<0), because we predicted their slopes would have opposite signs. Habitat-specific parasites (strong frequency bias) are indicated by filled points. Habitat-specific alleles are indicated by larger points. We combine all three lake-stream pairs in this plot, because different alleles were involved in parasite susceptibility or resistance in each pair.

**Figure 8.**
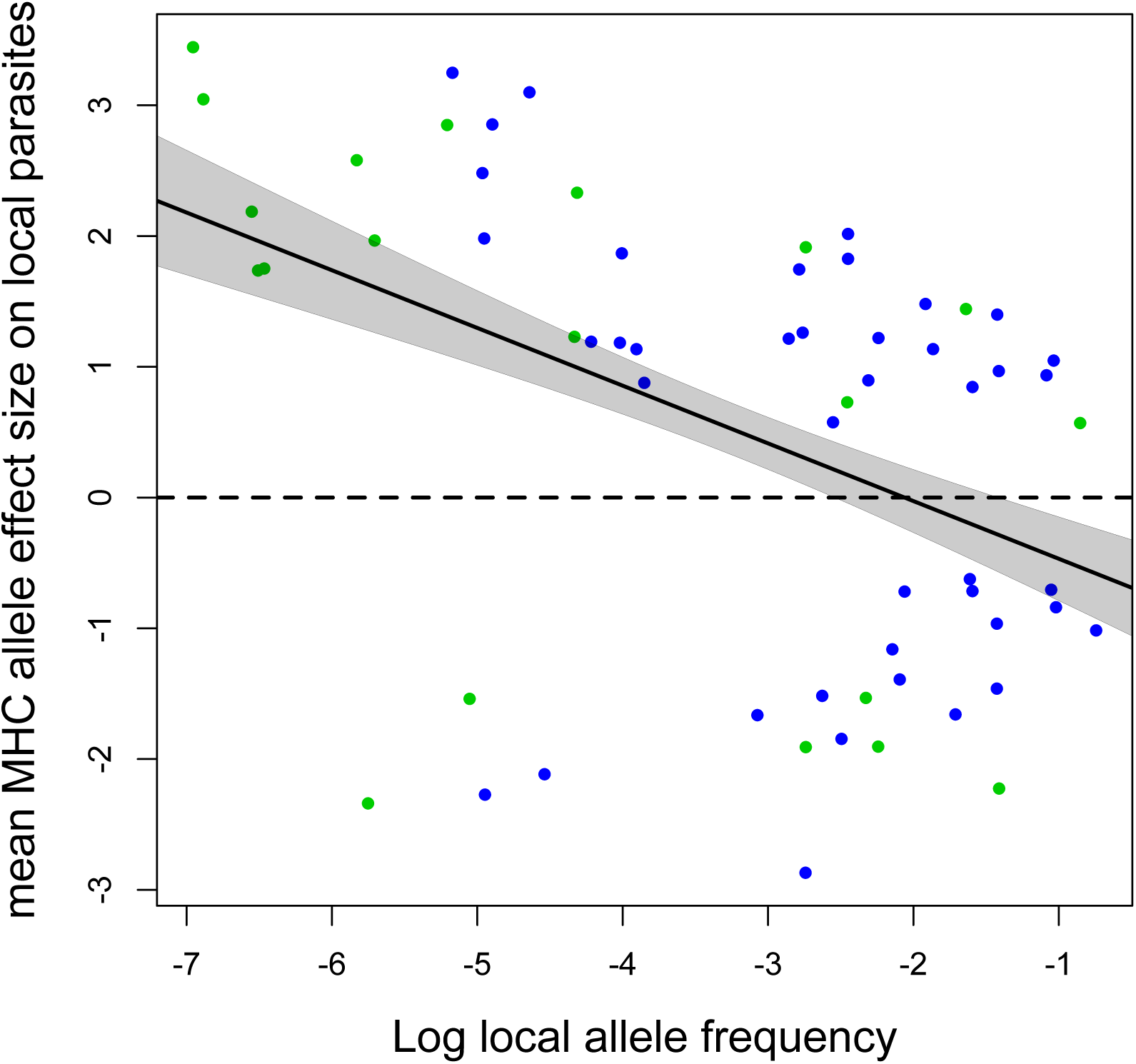
Locally rare alleles tend, on average, to confer susceptibility to local parasites (positive allele-parasite associations), whereas common alleles tend to be about equally likely to confer resistance or susceptibility. Allele frequency is calculated as the log_2_ of the fraction of individuals carrying the allele. Each point is an allele in a particular habitat in a lake-stream pair. For each allele, we calculated its average effect size (β_m_.) across all present parasites. Only those alleles with at least some significant parasite associations are included.

## Discussion

Stickleback in lake and stream habitats harbor distinct but overlapping MHC class II genotypes, and substantial MHC diversity within populations (Chain *et al.* 2014; Eizaguirre *et al.* 2012a; Stutz & Bolnick 2014). We tested whether this between- and within-population MHC variation systematically supports a role of divergent versus balancing selection. We estimated pairwise statistical associations between 374 MHC alleles and 34 parasite taxa in three replicate lake-stream pairs, revealing a moderate number of clear associations between the presence of an MHC allele and less or greater infection by a particular parasite. This represents an exceptional survey of MHC-parasite associations in nature, revealing many more MHC-parasite associations than we expected, though most alleles were strongly associated with just a single parasite, if any. These associations included both positive and negative effects that may be loosely interpreted as evidence of susceptibility and resistance respectively.

These associations do not, by themselves, provide a clear test for divergent or balancing selection (Piertney & Oliver 2006). Instead, the MHC-parasite associations should be interpreted in the context of the alleles’ and parasites absolute or relative frequencies in each habitat (Fig. 1). To our knowledge, this has not previously been reported. We found a few MHC-parasite associations that clearly supported with either balancing or divergent selection. But, was no overall tendency for immigrant MHC alleles to be more susceptible to local parasites (implying divergent selection) or less susceptible (implying balancing selection). Instead, we found a weak but unexpected trend for locally rare (but not foreign) alleles to confer greater infection susceptibility.

We observed significant between-habitat differences in parasite community composition. A few parasites were systematically habitat-specific or habitat-biased (*Crepidostomum*, *Unionidae*, Thersitina, and Blackspot in lakes; Anisakis, Apatemon and Bunodera in streams). Consistent with prior findings from other lake-stream pairs (Eizaguirre *et al.* 2010; Feulner *et al.* 2015), stream parasite communities were a less diverse subset of the neighboring lake parasites.

We expected these differences in parasite community composition to generate divergent natural selection on stickleback immune genes. Consistent with this expectation, we observed significant differences in MHC genotypes between parapatric lake and stream sites. Approximately 10% of alleles exhibit substantial (> 2-fold) frequency differences between parapatric habitats. But,genomic SNPs (most of which will be neutral) also exhibit significant divergence along the sampled clines (Weber *et al.* 2017). Despite divergence in MHC genotypes, we saw little evidence for local adaptation. A few alleles are rare in habitats where they confer susceptibility to a parasite (e.g., P273 is rare in Comida Lake where Unionidae is common). Such patterns may arise if selection largely eliminates alleles from habitats where they are detrimental. However, this depletion of locally susceptible alleles involves only a few MHC alleles from Comida. Taking a larger view across many allele-parasite associations, we found no general trend for alleles common in a given habitat to confer (i) protection against that habitat’s parasites, or (ii) susceptibility to parasites in the neighboring habitat. The prediction illustrated in Fig. 1B is therefore not supported.

In a few cases, we found rare MHC alleles associated with reduced infection rates. This apparent rare-allele advantage fits expectations from balancing selection. However, as with local adaptation, this particular outcome is confined to a few examples. There is no overall tendency for locally rare alleles to be more resistant (or, locally common alleles to be more susceptible). We therefore found no overall support for balancing selection in any of the three lake-stream pairs, despite trying multiple variants on our analytical approach.

An alternative approach to testing balancing selection is to ask whether MHC-parasite associations depend on allele frequency *within* a given habitat, rather than frequency differences between habitats. We find that locally rare alleles tend to confer susceptibility to local parasites, whereas locally common alleles are equally likely to be susceptible or resistant. This result is consistent with the notion that selection removes alleles that are susceptible to local parasites. But, by focusing specifically on local allele frequency (rather than between-habitat frequency difference), this result does not prove that *different* alleles are favored in the two habitats. Indeed, the susceptible rare alleles are typically not immigrants (e.g., not more common in the other habitat).

To summarize, we proposed contrasting predictions to distinguish between balancing versus divergent selection. Both predictions were supported by a few allele-parasite combinations, but neither was supported overall. There may be several explanations for these equivocal results. First, divergent- and balancing selection may in fact be weak or absent. This conclusion would be odd, given the high MHC diversity within populations and divergence between adjoining populations that readily exchange migrants. Second, MHC has been widely linked to mate choice decisions in stickleback and other vertebrates (Lenz *et al.* 2009; Milinski 2006), so divergent sexual selection, might plays a primary role in MHC population structure. Lastly, divergent and balancing selection might act concurrently. As we show here, some parasites might drive divergence in some alleles’ frequencies, while other parasites target locally common alleles. But, the net effect may be that these two selective processes obscure each other’s signals in an overall meta-analysis, as we find.

Some additional caveats are worth noting. Our results are based on a brief survey of three lake-stream pairs in a single season and year. It may be that the strongest selection occurs at another season, ontogenetic stage, or year. Also, we focused exclusively on readily visible macroparasites, but MHC evolution could plausibly also depend on readily overlooked symbionts including but not limited to gut microbiota (Bolnick *et al.* 2014). Lastly, although some stickleback parasites are well known to reduce host fitness, we do not presently know how host survival or fecundity depend on infection loads of all parasites examined here.

We observed a substantial number of MHC allele –parasite associations, consistent with typical expectations that MHC IIβ is involved in immunity to macroparasites. However, it is surprising that positive and negative associations (‘susceptibility’ and ‘resistance’, respectively) were about equally common. Why would so many MHC alleles, when present in a fish, coincide with greater infection by a certain parasite? A first possibility is that positive effects are spurious consequences of having alternative alleles. The presence of one allele may imply the absence of an alternative allele with protective value. This explanation is unlikely in our present study, because we statistically accounted for moderately correlated alleles. A second explanation could be that an allele facilitating recognition of one parasite might result in immunological trade-offs that inhibit resistance to another parasite. For instance, an MHC allele that recognized a microbe might drive an inflammatory response that inhibits resistance to a subsequent helminth infection (Moser & Murphy 2000; Oladiran & Belosevic 2012; Salgame *et al.* 2013). A third possibility entails direct interactions among parasites. If one parasite inhibits invasion of another parasite, then an allele that resists the former may facilitate infection by the latter (Hafer & Milinski 2015). Lastly, because we are sampling wild-caught adult fish, a positive correlation between genotype and infection could reflect a tolerance effect of the allele. If individuals with a given allele are more likely to survive a chronic infection, the allele will be enriched among infected survivors, compared to uninfected individuals(Westerdahl *et al.* 2012). This last point brings up an important caveat about our analysis: we assume that higher infection load implies lower fitness, but variation in infection tolerance, and survival prior to our sampling effort, complicates this interpretation.

Prior studies of stickleback have suggested that MHC heterozygosity is itself under stabilizing selection (Wegner *et al.* 2004; Wegner *et al.* 2003). The suggestion is that individuals with few MHC alleles are unable to recognize enough parasites, whereas individuals with too many alleles have reduced T-cell receptor diversity, resulting in an intermediate optimal MHC heterozygosity. Prior studies suggested that lower parasite diversity in streams than in lakes, causes a lower optimal allelic diversity (Feulner *et al.* 2015). In contrast, we find comparable MHC diversity in the lake and stream populations, even though the stream parasite community is less diverse. Moreover, stream sticklebacks had lower effective population sizes based on genomic SNP data (Weber et al. 2017). So, MHC diversity is actually relatively high in the stream (compared to neutral markers) despite their lower parasite diversity. Our samples therefore do not support the proposal that stream stickleback have a lower optimal MHC diversity.

## Conclusions

A great many studies have tested for divergent or balancing selection on MHC, in numerous vertebrate species (reviewed by (Bernatchez & Landry 2003; Edwards & Hedrick 1998; Eizaguirre & Lenz 2010; Hedrick 2002; Piertney & Oliver 2006; Yasukochi & Satta 2013). Few of these studies have simultaneously tested for both forms of selection (Tobler *et al.* 2014). Most of these studies yield some support for one hypothesis or the other, but frequently the supporting evidence has important caveats and some inconsistencies. Consequently, the evolutionary maintenance of MHC diversity within and between populations remains something of a puzzle despite extensive research. Our own data exacerbate this puzzle. We found some support for both divergent and balancing selection, depending on which allele and parasite we considered. But, at the scale of all alleles and parasites, we found no predominant signal favoring one form of selection over the (Fig. 1).

We propose that there in fact may not be a predominant form of selection at this multi-locus gene family. Rather, balancing and divergent selection act simultaneously on different MHC II alleles, in association with different parasites. Some alleles may experience a native advantage, while others may experience a rare-allele advantage. Current analytical approaches are not effective at separating such simultaneous forms of selection. Future work on MHC evolution must therefore account for many parasite species concurrently, and the distinct but simultaneous selective pressures that each may exert.

## Acknowledgments

We thank Claire Patenia and Jason Clu for lab assistance. This research was supported by fellowships to DIB from the David and Lucile Packard Foundation and the Howard Hughes Medical Institute, and a grant to WES from the Graduate Program in Ecology Evolution and Behavior at UT Austin.

## Data Accessibility

Data required to reproduce the analyses presented in this paper will be made publically available at the time of publication. 454 amplicon sequencing reads have been uploaded to____. Tables of allele presence/absence from these 454 amplicon sequences will be uploaded to Dryad doi:____. Tables of parasite infections and fish ecomorphology and sex will be uploaded to Dryad at doi:____, along with meta-data linking individual fish to their MHC genotype and sampling location.

## Author contributions

WES and DIB collaboratively planned the data collection and analysis. WES conducted the field work, lab work, sequencing, bioinformatics. The statistical analyses were conducted primarily by WES with contributions from DIB. WES and DIB wrote the manuscript together.

## SUPPORTING INFORMATION

**Table S1.** Sampling locations and sample sizes. Columns give distances downstream from lake (in meters) where fish were sampled. Cells indicate the total number of fish sampled at each sampling distance. Not all fish were screened for parasites or genotyped.

|  | Lake location | Stream sites (distance downstream from lake, m) |  |  |  |  |  |  |  |  |  |  |  |  |  |
| --- | --- | --- | --- | --- | --- | --- | --- | --- | --- | --- | --- | --- | --- | --- | --- |
| Pair | Latitude/Longitude | Lake | 20 | 25 | 50 | 100 | 150 | 200 | 250 | 300 | 400 | 500 | 1000 | 1500 | stream total |
| Comida | 50.1443, -125.5283 | 81 | 0 | 0 | 14 | 20 | 21 | 20 | 20 | 19 | 20 | 20 | 20 | 20 | 194 |
| Farewell | 50.2010, -125.5860 | 126 | 0 | 0 | 0 | 13 | 0 | 50 | 0 | 48 | 50 | 50 | 50 | 0 | 261 |
| Roberts | 50.2266, -125.5530 | 138 | 1 | 1 | 0 | 43 | 4 | 48 | 47 | 49 | 28 | 18 | 50 | 70 | 359 |

**Table S2.** sample sizes by data available. Cells indicate the total number of fish at each site for which each type data was collected (dissected=measured and screened for parasites, genotyped=genotyped at MHC loci).

|  | Parasite screening | MHC genotyping | both |
| --- | --- | --- | --- |
| Comida Lake | 81 | 42 | 42 |
| Comida Stream | 190 | 138 | 136 |
| Farewell Lake | 121 | 102 | 97 |
| Farewell Stream | 258 | 211 | 209 |
| Roberts Lake | 137 | 119 | 118 |
| Roberts Stream | 333 | 234 | 216 |

**Table S3.**
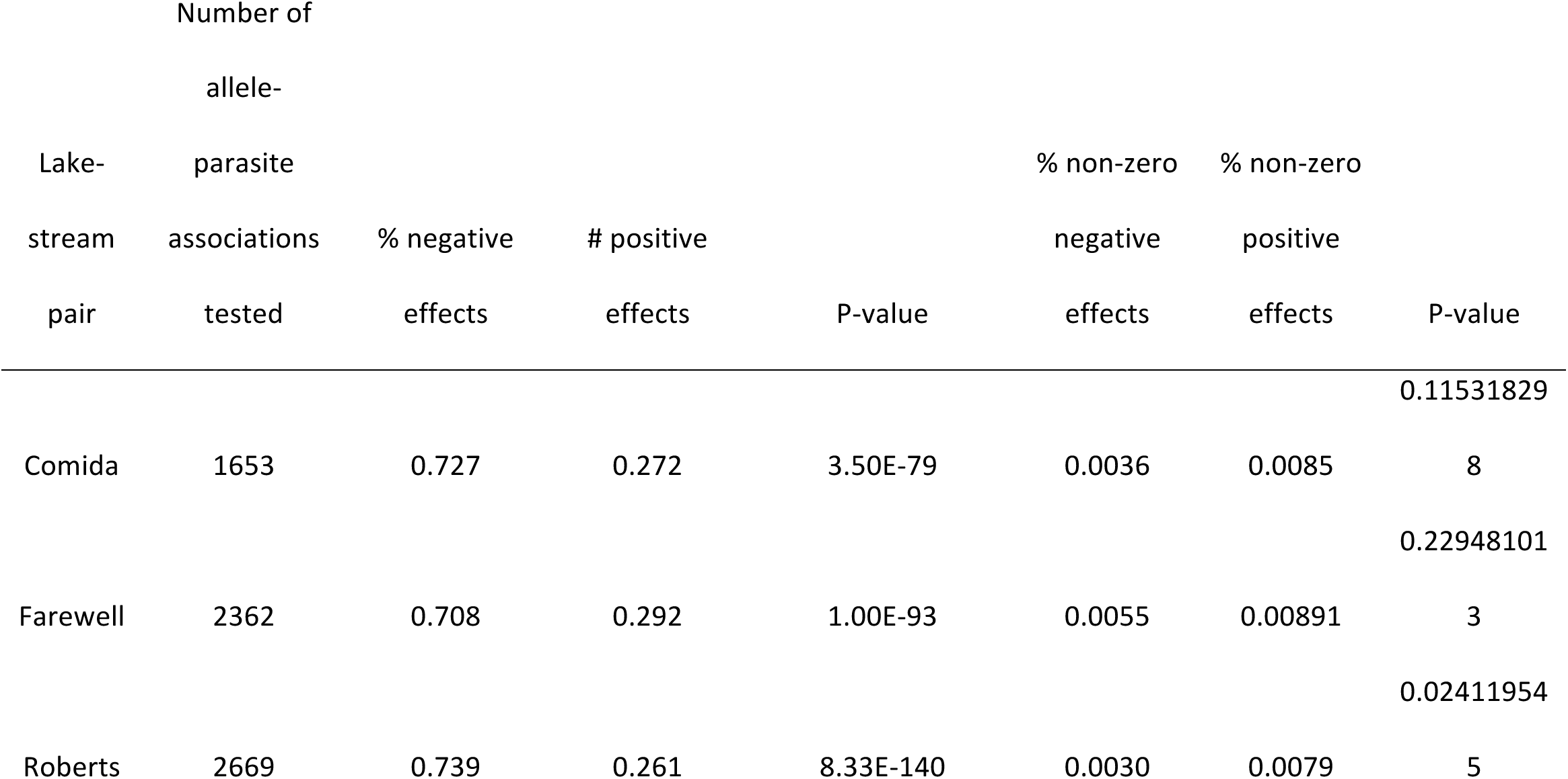
Percentage of estimated allele effects that were positive and negative. Numbers in parentheses give the percentage of estimated effects that were non-zero effects. P-values are derived from exact binomial tests that negative and positive effects were equally likely within each pair ( p-value in parenthesis gives the result for tests for just non-zero effects).

**Table S4.** Estimated non-zero effects. Habitat specific parasite and alleles are indicated in bold.

| pair | allele m | parasite p | $\beta_{mp}$ | 95% HPDI | | Habitat Specificity | |
| --- | --- | --- | --- | --- | --- | --- | --- |
|  |  |  |  | low | hi | parasite | allele |
| Comida | P293 | Crepidostomum | -2.87 | -5.17 | -0.78 | **lake** | none |
| Farewell | P198 | Crepidostomum | -2.27 | -4.88 | -0.13 | **lake** | none |
| Comida | P70 | Apatemon | -2.34 | -4.67 | -0.13 | **stream** | none |
| Roberts | P236 | Cestode_spp2 | -1.91 | -4.42 | -0.03 | none | none |
| Roberts | P115 | Anisakis | -2.23 | -4.13 | -0.56 | none | none |
| Farewell | P46 | Eustrongyloides | -1.91 | -3.96 | -0.02 | none | none |
| Roberts | P163 | Capillaria | -2.12 | -3.92 | -0.23 | none | none |
| Farewell | P139 | Unident_spine_cysts | -1.85 | -3.76 | -0.08 | **lake** | none |
| Farewell | P150 | Crepidostomum | -1.66 | -3.25 | -0.19 | **lake** | none |
| Farewell | P182 | Capillaria | -1.66 | -3.17 | -0.11 | none | none |
| Farewell | P395 | Eustrongyloides | -1.53 | -3.09 | -0.02 | none | none |
| Comida | P269 | Apatemon | -1.54 | -2.98 | -0.15 | **stream** | none |
| Farewell | P403 | Capillaria | -1.39 | -2.94 | -0.16 | none | none |
| Roberts | P32 | Capillaria | -1.46 | -2.94 | -0.11 | none | **lake** |
| Farewell | P163 | Crepidostomum | -1.52 | -2.63 | -0.22 | **lake** | none |
| Comida | P299 | Diplostomum indistinctum | -1.16 | -2.25 | -0.01 | **lake** | none |
| Roberts | P216 | Cestode_spp5 | -1.02 | -1.97 | -0.24 | **lake** | none |
| Roberts | P32 | Blackspot | -0.96 | -1.87 | -0.04 | none | **lake** |
| Farewell | P46 | Apatemon | -0.72 | -1.51 | 0.00 | none | none |
| Comida | P393 | Crepidostomum | -0.71 | -1.46 | -0.03 | **lake** | none |
| Roberts | P1 | Blackspot | -0.72 | -1.41 | -0.04 | none | none |
| Comida | P267 | Thersitina | -0.84 | -1.40 | -0.29 | none | none |
| Farewell | P397 | Crepidostomum | -0.62 | -1.22 | -0.01 | **lake** | none |
| Comida | P292 | Bunodera | 0.73 | 0.01 | 1.37 | none | none |
| Farewell | P23 | Nematode_spp7 | 1.44 | 0.02 | 2.93 | none | none |
| Roberts | P1 | Cestode_spp5 | 0.85 | 0.02 | 1.88 | **lake** | none |
| Comida | P272 | Diplostomum indistinctum | 0.88 | 0.02 | 1.75 | **lake** | none |
| Farewell | P103 | Apatemon | 1.14 | 0.02 | 2.47 | none | **lake** |
| Comida | P265 | Bunodera | 0.57 | 0.02 | 1.17 | none | none |
| Roberts | P341 | Blackspot | 1.19 | 0.03 | 2.21 | none | none |
| Comida | P273 | Unionidae | 1.18 | 0.03 | 2.58 | **lake** | **stream** |
| Comida | P59 | Apatemon | 1.74 | 0.04 | 3.54 | **stream** | **lake** |
| Farewell | P412 | Thersitina | 1.14 | 0.04 | 2.14 | **lake** | none |
| Roberts | P20 | Thersitina | 1.22 | 0.04 | 2.48 | **lake** | none |
| Roberts | P35 | Nematode_spp4 | 1.83 | 0.05 | 3.48 | none | none |
| Comida | P279 | Thersitina | 0.58 | 0.05 | 1.11 | none | none |
| Comida | P403 | Crepidostomum | 1.22 | 0.06 | 2.52 | **lake** | **lake** |
| Roberts | P8 | Capillaria | 0.94 | 0.07 | 1.77 | none | none |
| Roberts | P236 | Nematode_spp3 | 1.48 | 0.08 | 2.85 | none | none |
| Farewell | P82 | Blackspot | 2.48 | 0.09 | 5.07 | **lake** | none |
| Roberts | P376 | Blackspot | 0.90 | 0.09 | 1.79 | none | none |
| Farewell | P60 | Nematode_spp7 | 2.58 | 0.09 | 4.84 | none | none |
| Roberts | P110 | Cestode_spp5 | 0.97 | 0.10 | 1.87 | **lake** | none |
| Farewell | P218 | Eustrongyloides | 1.23 | 0.10 | 2.24 | none | none |
| Comida | P92 | Apatemon | 1.75 | 0.10 | 3.46 | **stream** | **lake** |
| Roberts | P231 | Cestode_spp5 | 1.26 | 0.10 | 2.67 | **lake** | **lake** |
| Farewell | P166 | Crepidostomum | 1.98 | 0.15 | 3.98 | **lake** | none |
| Roberts | P35 | Unionidae | 2.02 | 0.16 | 4.02 | **lake** | none |
| Roberts | P115 | Cestode_spp5 | 1.05 | 0.16 | 1.88 | **lake** | none |
| Farewell | P414 | Unident_spine_cysts | 1.40 | 0.17 | 2.55 | **lake** | none |
| Comida | P66 | Apatemon | 2.19 | 0.30 | 4.22 | **stream** | none |
| Farewell | P37 | Blackspot | 2.85 | 0.34 | 4.86 | **lake** | none |
| Farewell | P240 | Thersitina | 1.74 | 0.36 | 3.02 | **lake** | none |
| Farewell | P46 | Bunodera | 1.91 | 0.37 | 3.62 | none | none |
| Farewell | P91 | Schistocephalus | 1.87 | 0.39 | 3.23 | none | none |
| Farewell | P158 | Bunodera | 2.85 | 0.47 | 5.04 | none | none |
| Roberts | P190 | Cestode_spp2 | 2.33 | 0.51 | 3.91 | none | none |
| Comida | P350 | Apatemon | 1.97 | 0.53 | 3.52 | **stream** | none |
| Roberts | P97 | Cestode_spp7 | 3.10 | 0.67 | 5.53 | none | none |
| Roberts | P123 | Cestode_spp2 | 3.05 | 0.78 | 5.07 | none | none |
| Roberts | P155 | Cestode_spp2 | 3.44 | 1.17 | 5.62 | none | none |
| Roberts | P18 | Cestode_spp6 | 3.25 | 1.22 | 5.40 | none | none |

**Fig. S1.**
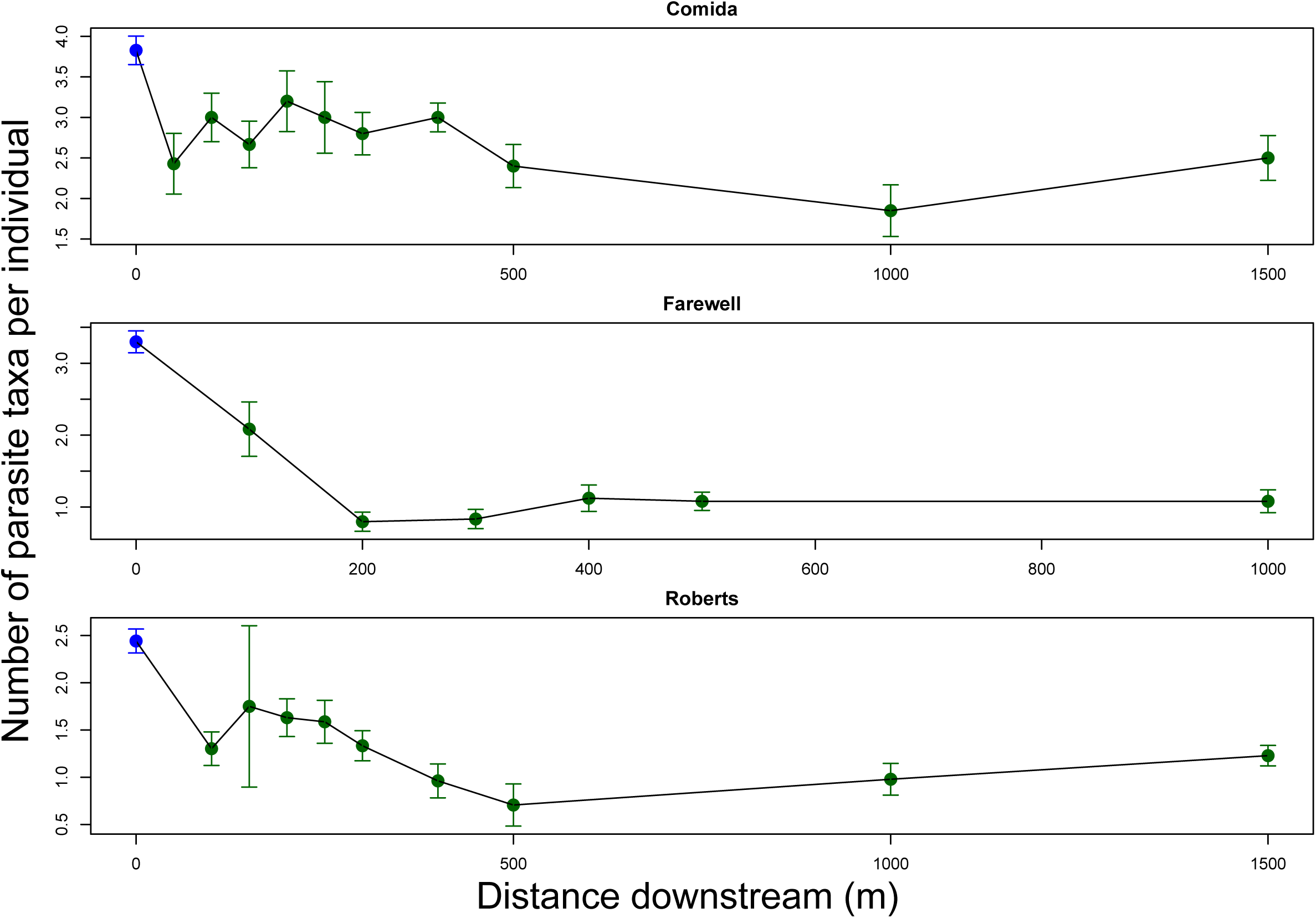
In all three lake-stream clines, stream stickleback carried a lower parasite richness than did their neighboring stream stickleback (all P<0.0001), with no significant effect of distance within the stream (all P>0.05).

**Figure S2.**
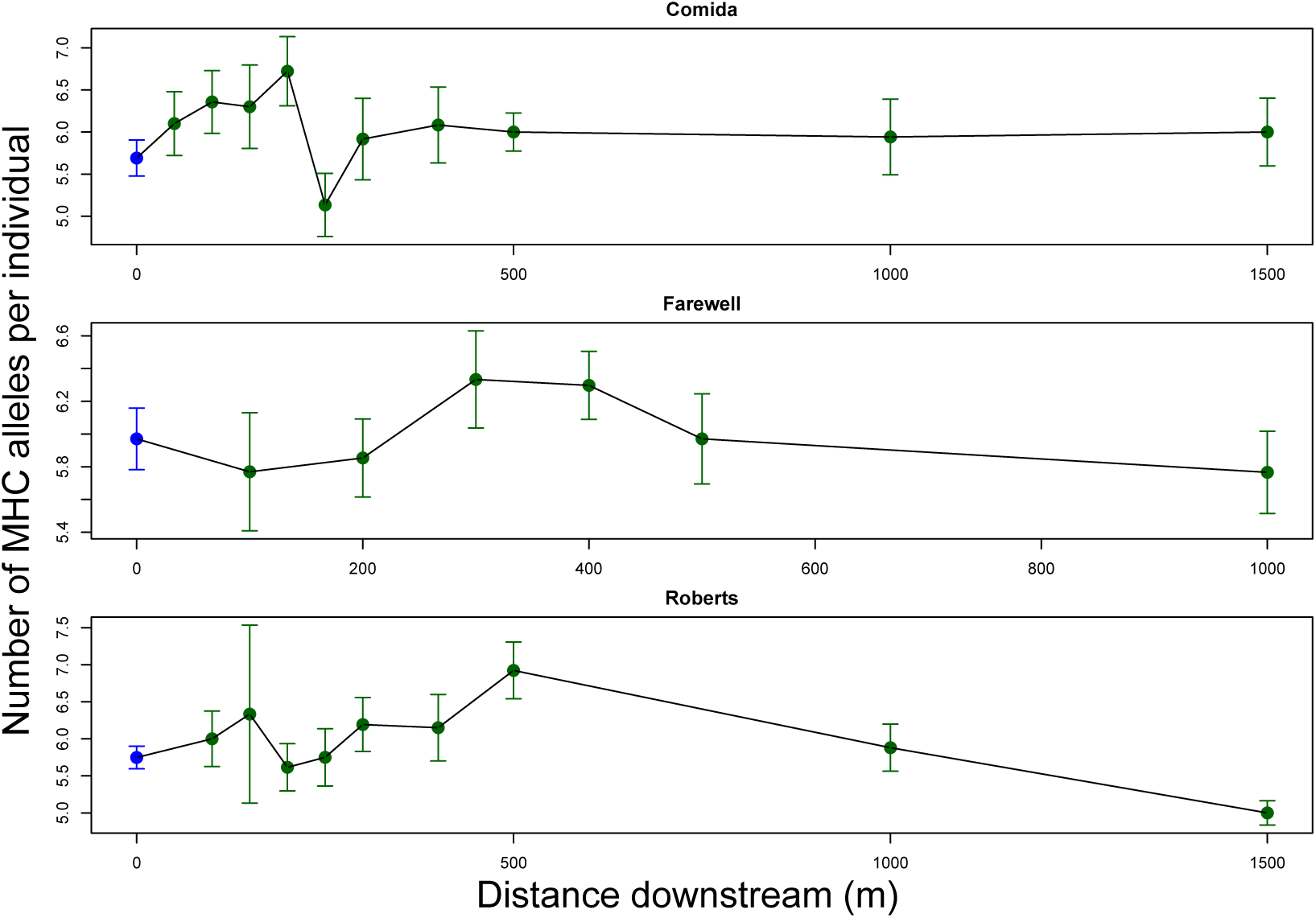
MHC allelic diversity (operationally defined here as the number of unique amino acid sequences per individual fish) varies across lake-stream clines. In the Comida and Farewell pairs, there is no significant effect of habitat, distance, nor a quadratic distance effect (ANCOVA; Comida: P=0.172, 0.433, and 0.785 respectively; Farewell P=0.832, 0.724, 0.128). In Roberts Lake, however, there is a significant effect of distance, including both a linear and a negative quadratic trend (P=0.0065 and 0.0201) but no effect of habitat (P=0.9373). In all three pairs, the greatest per-fish MHC diversity occurs in the stream, midway along the transect. This transitory increase in diversity is consistent with the proposed diversity-increasing effect of migration. Combining the three lake-stream pairs into a single ANOVA analysis, we find a significant effect of distance (P=0.0006) and marginal quadratic effect (P=0.091) on MHC diversity, but no effect of pair (P=0.118) or habitat (P=0.497). This result stands in contrast to other stickleback lake-stream pairs, in which stream fish consistently exhibit lower MHC diversity than lake fish (Eizaguirre *et al.* 2012a; Eizaguirre *et al.* 2010; Feulner *et al.* 2015).

**Figure S3.**
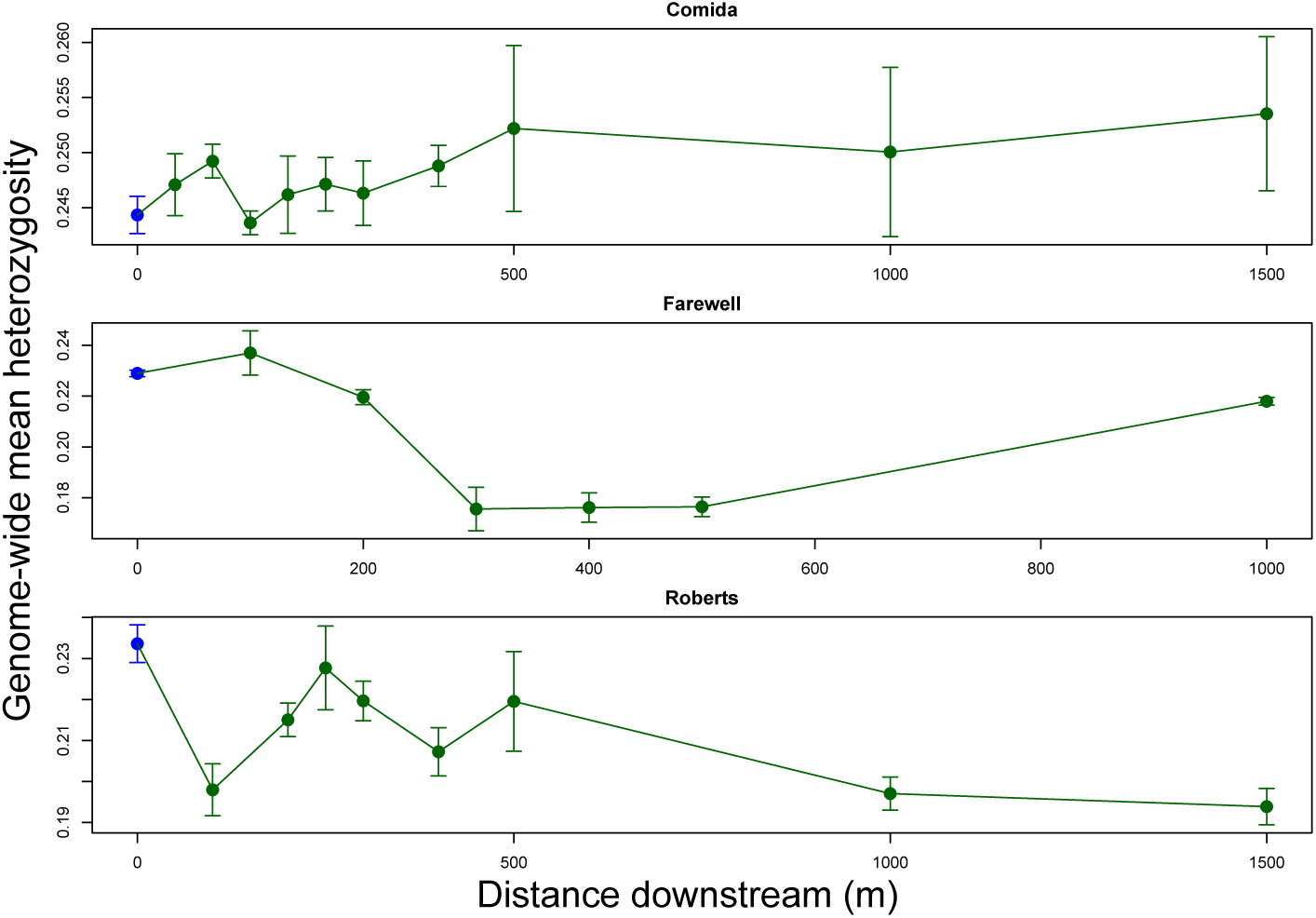
For comparison with Fig. S2, we here show the genomic SNP diversity (operationally defined here as the average heterozygosity across all scored SNPs) across lake-stream clines. In Comida Lake there is a weakly significant trend towards higher diversity farther downstream (P=0.034), and a very marginal difference between the habitats (P=0.095), but in an ANCOVA analysis with both effects, neither is significant. Farewell exhibits significant among-site variation in heterozygosity, driven by lower heterozygosity in the stream than in the lake (P<0.00001) and a quadratic effect of distance downstream (P<0.00001). Likewise, Roberts Lake exhibits lower heterozygosity in the stream than lake (P<0.00008) and decreasing heterozygosity with distance downstream (P=0.00315). Unlike the trend for MHC, genome-wide data shows no consistent tendency for genetic diversity to be elevated a short distance downstream. This suggests that the diversity-sustaining effect of migration might disproportionately impact MHC II sequences. That said, this comparison is only qualitative, because of the polygenic nature of MHC IIβ genotypes in this study.

**Fig. S4.**
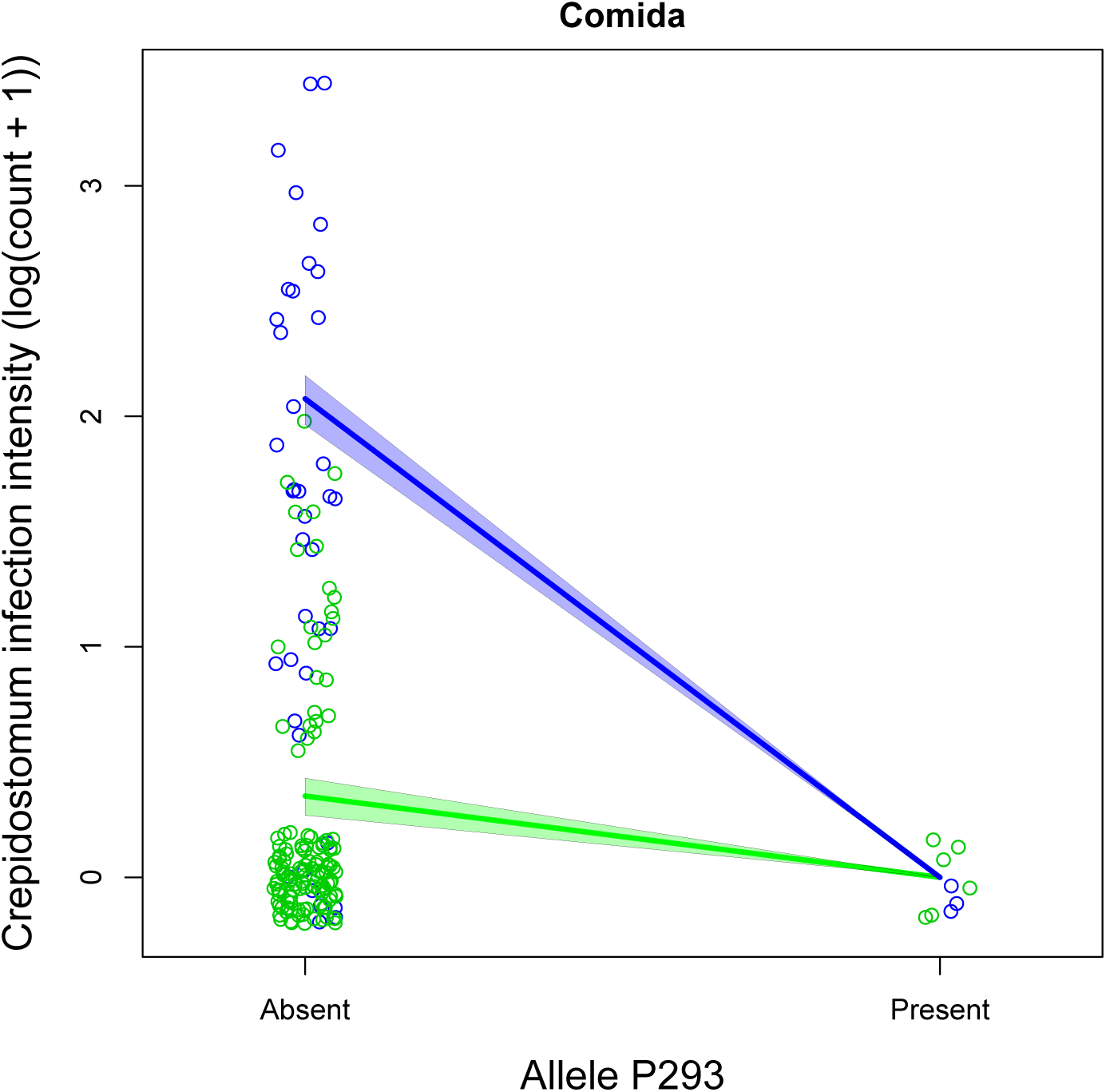
An example of a significant negative MHC allele-parasite association. Crepidostomum sp. infection intensity is lower in Comida fish carrying allele P293, than in fish without. This trend is supported overall, but is stronger for lake fish (shown in blue) than stream fish (green).

**Fig. S5.**
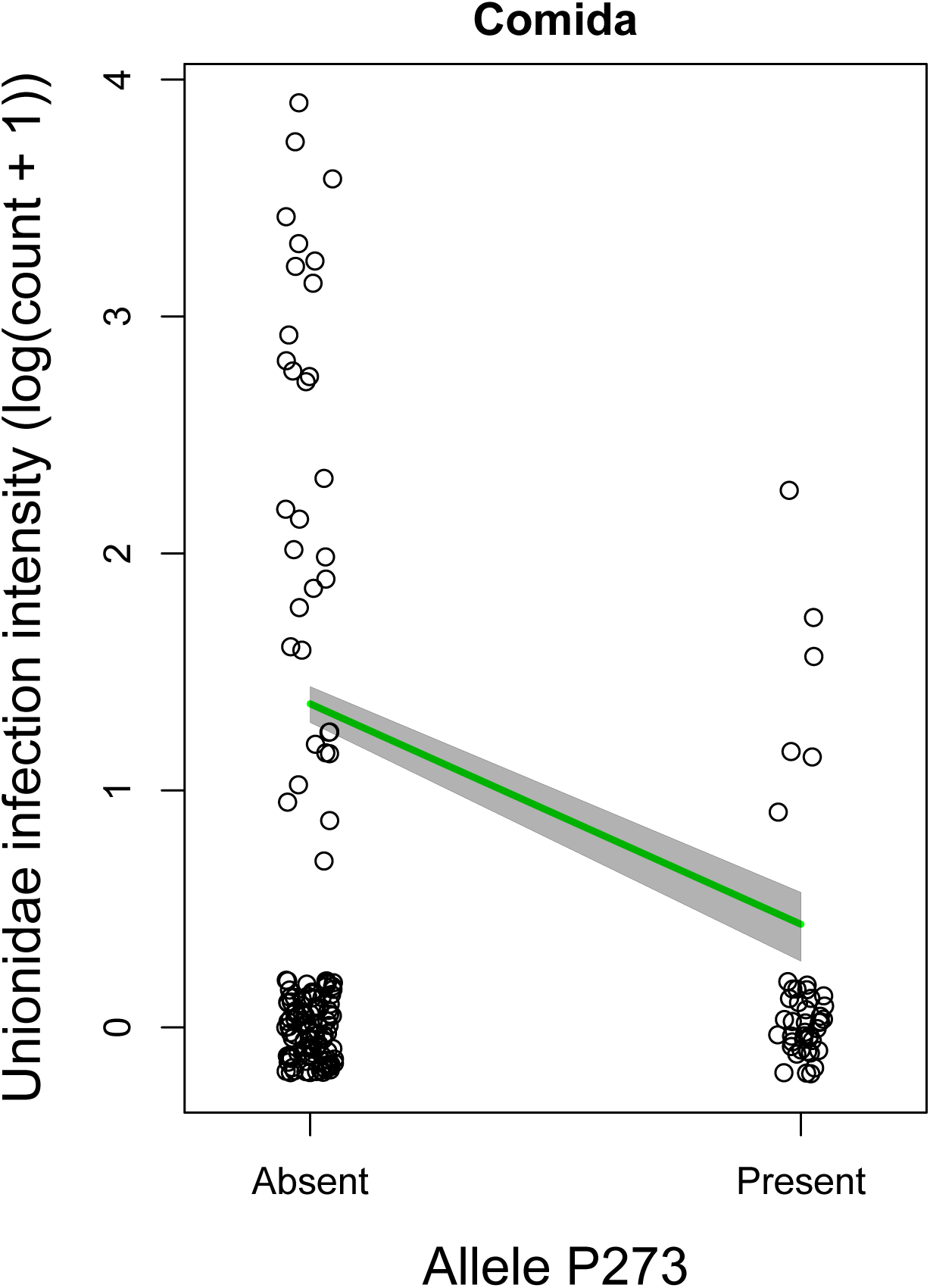
An example of a significant negative MHC allele-parasite association. Unionidae infection intensity is lower in Comida fish carrying allele P273, than in fish without. This trend is independent of fish habitat.

**Fig. S6.**
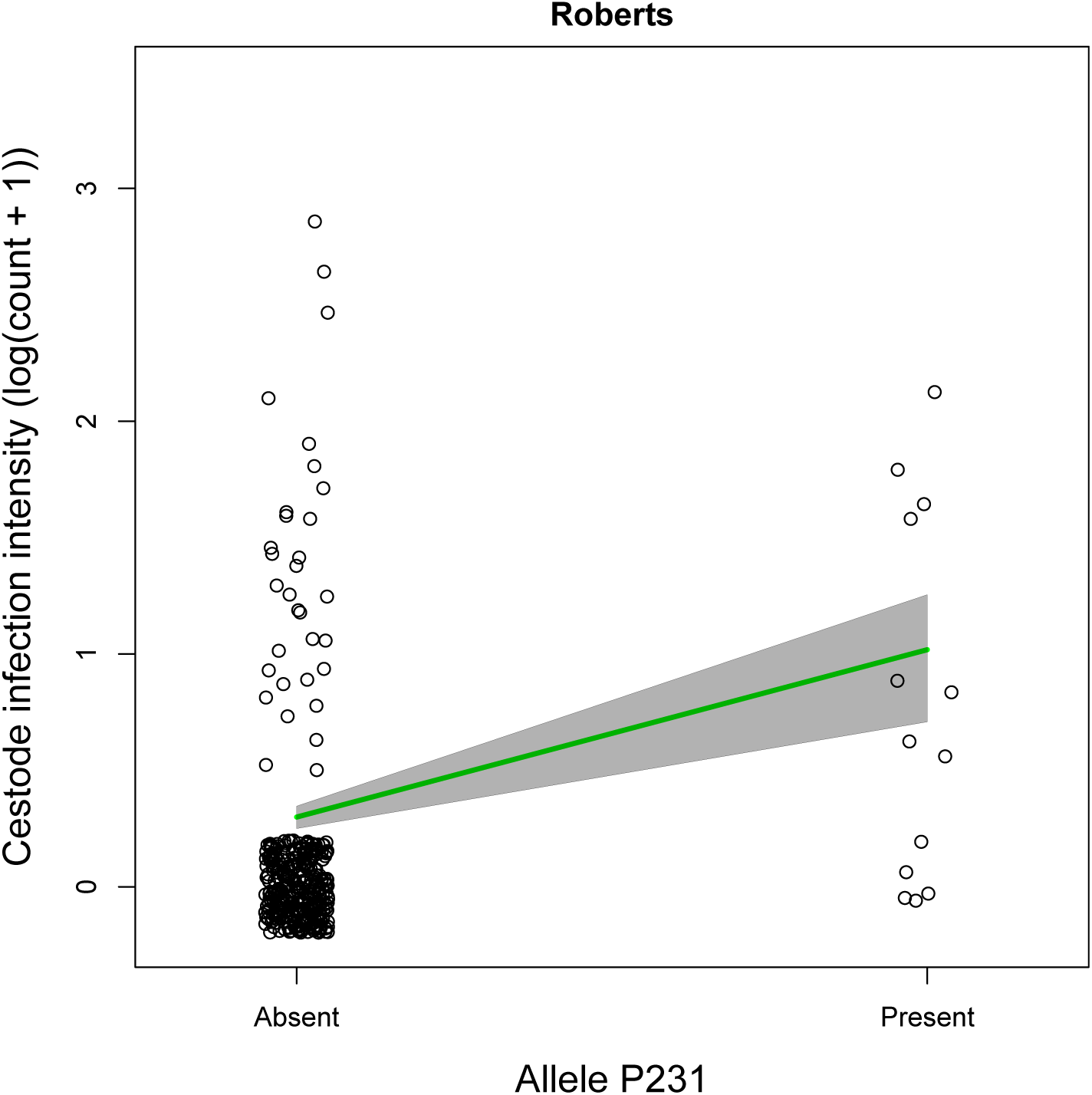
An example of a significant positive MHC allele-parasite association. Cestode spp infection intensity is higher in Roberts fish carrying allele P231, than in fish without.

**Fig. S7.**
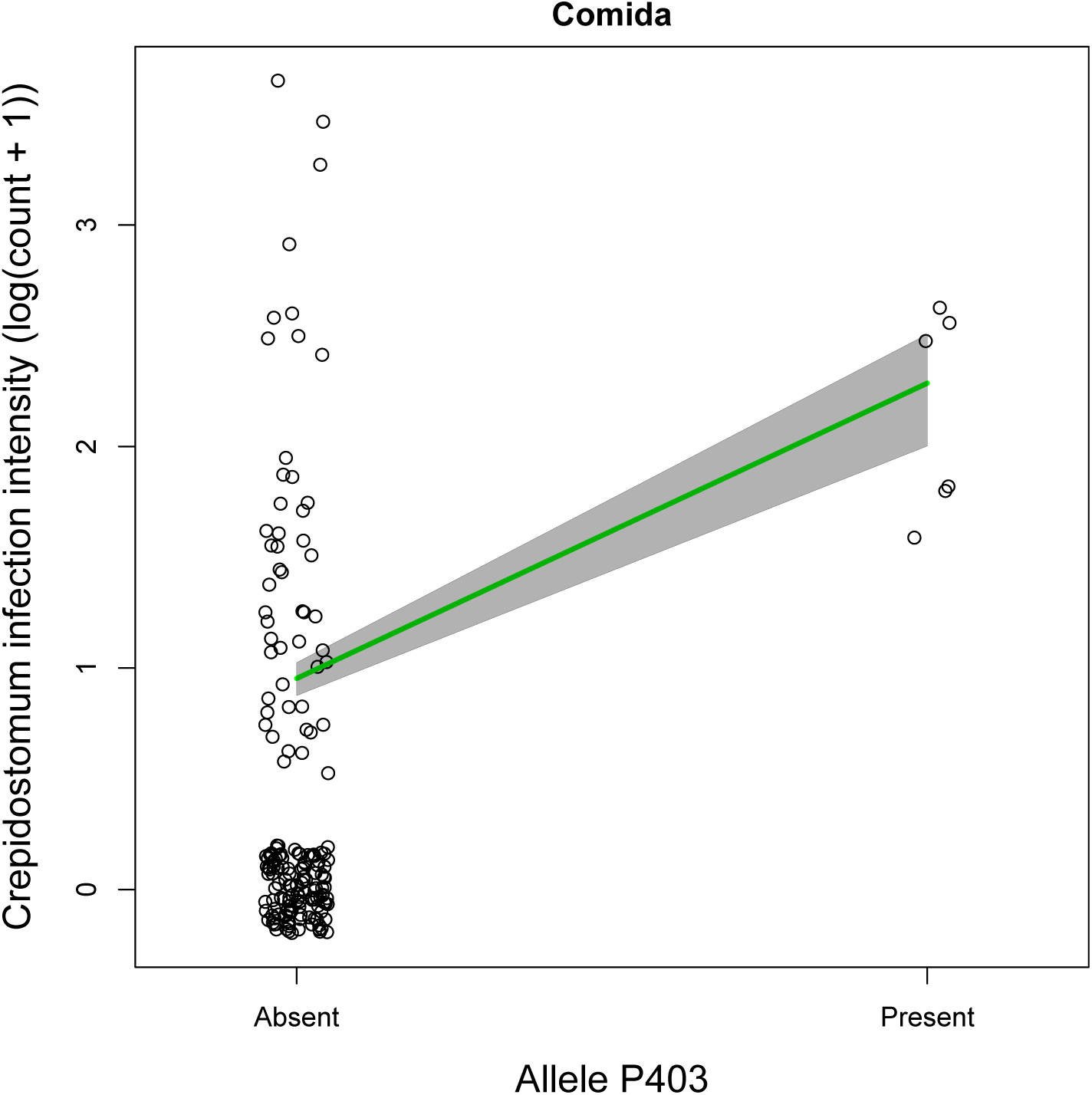
An example of a significant positive MHC allele-parasite association. Crepidostomum infection intensity is higher in Comida fish carrying allele P403, than in fish without. This trend is largely independent of the effect plotted in Fig. S4 for the same parasite and the same lake-stream pair, because the allele P403 shown here segregates independently of allele P293.

**Figure S8.**
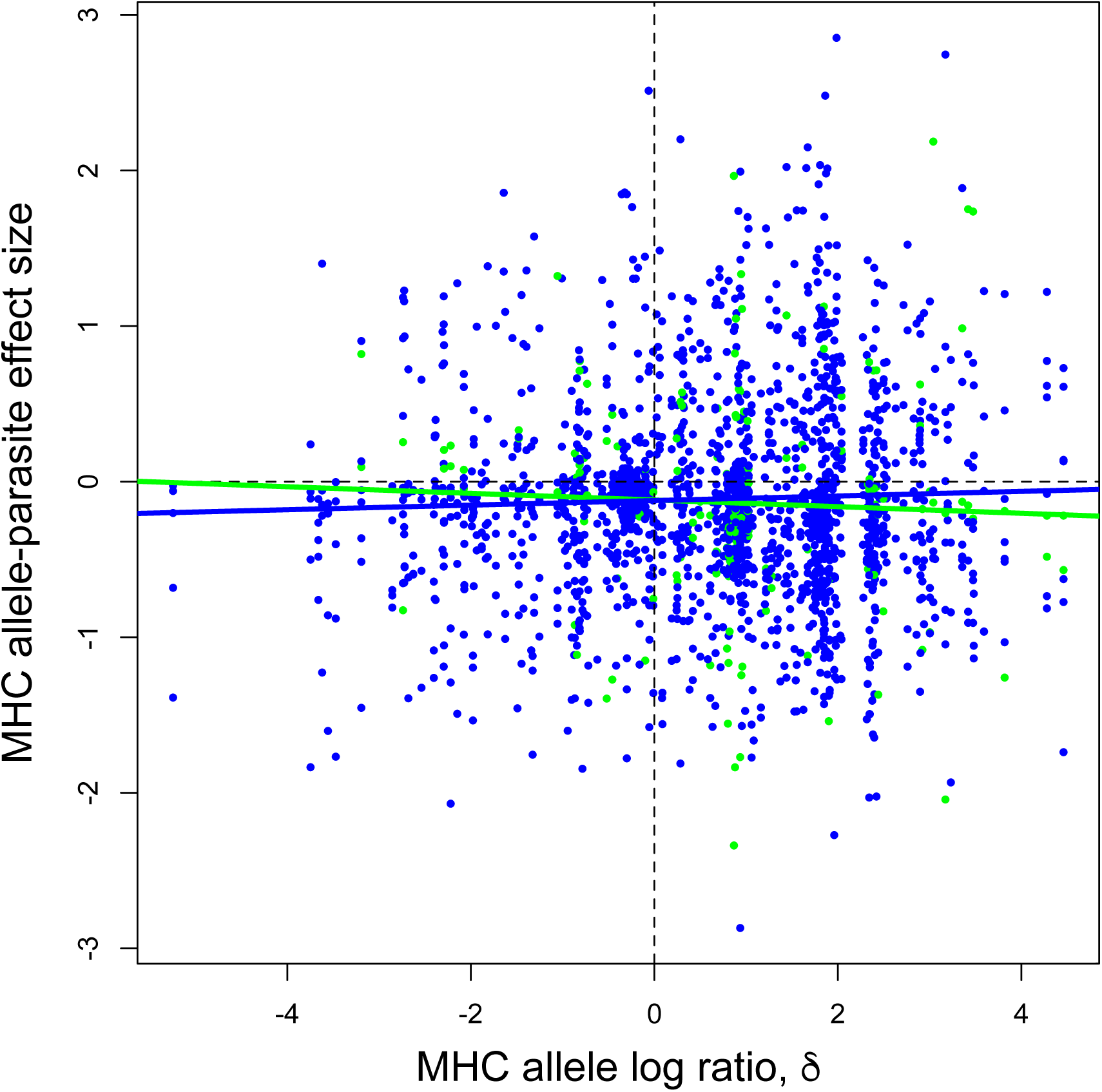
The same plot as in Fig. 7 in the main text, but for all allele-parasite associations tested, rather than just the strongly habitat-biased parasites, or strongly habitat-biased alleles.

## Supplemental Methods

### Accounting for allele co-occurrence

Given that MHC II-B exists as multiple paralog loci, alleles can be co-inherited as haplotype blocks, resulting in linkage disequilibrium between different alleles. Because alleles may be co-inherited, it could be difficult to determine which of two (or more) linked alleles are responsible for variation in infections. We therefore calculated Yule's Q (Warrens 2008) as a measure of association between all pairs of alleles within a lake/stream pair, using *Yule* function in the *psych* package in R 3.2.1 (Revelle 2016). Like Pearson correlation coefficients, Q ranges from −1 to 1 and indicates the degree of positive or negative association between two binary variables (e.g. allele presence). This metric was used to determine which alleles co-occurred strongly.

### Yule's Q results

Of the 27,406 pairwise comparisons of alleles, only 1702 pairs of alleles (6%) had a value of Yule's Q greater than 0.8, implying strong linkage. Many of these are cases where one allele is very rare (i.e. occurs once) and thus overlaps completely with any other alleles found in that particular individual. In general, the relatively low levels of co-occurrence means that allele effects could be estimated independently of the presence or absence of other alleles in most cases. There were 37 allele pairs that were reciprocally complete overlapping in occurrence; one allele from each of these pairs was removed for the data set prior to estimating effect sizes.

## References

Bernatchez L, Landry C (2003) MHC studies in nonmodel vertebrates: what have we learned about natural selection in 15 years? Journal of Evolutionary Biology 16, 363–377.

Berner D, Grandchamp A-C, Hendry AP (2009) Variable progress toward ecological speciation in parapatry: stickleback across eight lake-stream transitions. Evolution 63, 1740–1753.

Bolnick D, Snowberg LK, Caporaso JG, et al. (2014) Major Histocompatibility Complex IIB polymorphism contributes to among-individual variation in gut microbiota composition. Molecular Ecology 23, 4831–4845.

Chain FJ, Feulner PG, Panchal M, et al. (2014) Extensive copy-number variation of young genes across stickleback populations. PLoS Genet 10, e1004830.

Clague JJ, James TS (2002) History and isostatic effects of the last ice sheet in southern British Columbia. Quaternary Science Reviews 21, 71–87.

Copley R, Blais J, Rico C, et al. (2007) MHC adaptive divergence between closely related and sympatric african cichlids. PLoS ONE 2, e734.

Doherty PC, Zinkernagel RM (1975) Enhanced immunological surveillance in mice heterozygous at the H-2 gene complex. Nature 256, 50–52.

Edwards SV, Hedrick PW (1998) Evolution and ecology of MHC molecules: from genomics to sexual selection. Trends in Ecology & Evolution 13, 305–311.

Eizaguirre C, Lenz TL (2010) Major histocompatibility complex polymorphism: dynamics and consequences of parasite-mediated local adaptation in fishes. Journal of Fish Biology 77, 2023–2047.

Eizaguirre C, Lenz TL, Kalbe M, Milinski M (2012a) Divergent selection on locally adapted major histocompatibility complex immune genes experimentally proven in the field. Ecology Letters 15, 723–731.

Eizaguirre C, Lenz TL, Kalbe M, Milinski M (2012b) Rapid and adaptive evolution of MHC genes under parasite selection in experimental vertebrate populations. Nature communications 3, 621.

Eizaguirre C, Lenz TL, Sommerfeld RD, et al. (2010) Parasite diversity, patterns of MHC II variation and olfactory based mate choice in diverging three-spined stickleback ecotypes. Evolutionary Ecology 25, 605–622.

Evans ML, Neff BD, Heath DD (2010) MHC-mediated local adaptation in reciprocally translocated Chinook salmon. Conservation Genetics 11, 2333–2342.

Feulner PG, Chain FJ, Panchal M, et al. (2015) Genomics of divergence along a continuum of parapatric population differentiation. PLoS Genet 11, e1004966.

Figueroa F, Gúnther E, Klein J (1988) MHC polymorphism pre-dating speciation. Nature 335, 265–267.

Gandon S (2002) Local adaptation and the geometry of host-parasite coevolution. Ecology Letters 5, 246–256.

Gelman A, Hill J (2006) Data Analysis Using Regression and Multilevel/Hierarchical Models Cambridge University Press, New York.

Gelman A, Jakulin A, Pittau MG, Su Y-S (2008) A weakly informative default prior distribution for logistic and other regression models. Annals of Applied Statistics 2, 1360–1383.

Hadfield J (2010) MCMC methods for multi-response generalized linear mixed models: the MCMCglmm R package. Journal of Statistical Software 33, 1–22.

Hafer N, Milinski M (2015) When parasites disagree: evidence for parasite-induced sabotage of host manipulation. Evolution 69, 611–620.

Hedrick PW (2002) Pathogen resistance and genetic variation at MHC loci. Evolution 56, 1902–1908.

Hendry AP, Kaeuffer RE, Crispo E, Peichel CL, Bolnick DI (2013) Evolutionary inferences from exchangeability: individual classification approaches based on the ecology, morphology, and genetics of lake-stream stickleback population pairs. Evolution 67, 3429–3441.

Hill AV, Allsopp CE, Kwiatkowski D, et al. (1991) Common west African HLA antigens are associated with protection from severe malaria. Nature 352, 595–600.

Jager I, Eizaguirre C, Griffiths SW, et al. (2007) Individual MHC class I and MHC class IIB diversities are associated with male and female reproductive traits in the three-spined stickleback. Journal of Evolutionary Biology 20, 2005–2015.

Kalbe M, Eizaguirre C, Dankert I, et al. (2009) Lifetime reproductive success is maximized with optimal major histocompatibility complex diversity. Proceedings of the Royal Society of London B: Biological sciences 276, 925–934.

Kubinak JL, Stephens WZ, Soto R, et al. (2015) MHC variation sculpts individualized microbial communities that control susceptibility to enteric infection. Nature Communications 6, 8642.

Kurtz J, Kalbe M, Aeschlimann PB, et al. (2004) Major histocompatibility complex diversity influences parasite resistance and innate immunity in sticklebacks. Proceedings of the Royal Society B: Biological Sciences 271, 197–204.

Lamaze FC, Pavey SA, Normandeau E, et al. (2014) Neutral and selective processes shape MHC gene diversity and expression in stocked brook charr populations (Salvelinus fontinalis). Molecular Ecology 23, 1730–1748.

Lenz TL, Eizaguirre C, Scharsack JP, Kalbe M, Milinski M (2009) Disentangling the role of MHC-dependent ‘good genes’ and ‘compatible genes’ in mate-choice decisions of three-spined sticklebacks *Gasterosteus aculeatus* under semi-natural conditions. Journal of Fish Biology 75, 2122–2142.

Lohm J, Grahn M, Langefors A, et al. (2002) Experimental evidence for major histocompatibility complex-allele-specific resistance to a bacterial infection. Proceedings of the Royal Society B: Biological Sciences 269, 2029–2033.

Loiseau C, Richard M, Garnier S, et al. (2009) Diversifying selection on MHC class I in the house sparrow *(Passer domesticus)*. Molecular Ecology 18, 1331–1340.

Matthews B, Harmon LJ, M’Gonigle L, Marchinko KB, Schaschl H (2010) Sympatric and allopatric divergence of MHC genes in threespine stickleback. PLoS ONE 5, e10948.

McCairns RJS, Bourget S, Bernatchez L (2011) Putative causes and consequences of MHC variation within and between locally adapted stickleback demes. Molecular Ecology 20, 486–502.

Meyer D, Thomson G (2001) How selection shapes variation of the human major histocompatibility complex: a review. Annals of Human Genetics 65, 1–26.

Milinski M (2006) The Major Histocompatibility Complex, sexual selection, and mate choice. Annual Review of Ecology, Evolution, and Systematics 37, 159–186.

Miller HC, Allendorf F, Daugherty CH (2010) Genetic diversity and differentiation at MHC genes in island populations of tuatara (Sphenodon spp.). Molecular Ecology 19, 3894–3908.

Mona S, Crestanello B, Bankhead-Dronnet S, et al. (2008) Disentangling the effects of recombination, selection, and demography on the genetic variation at a major histocompatibility complex class II gene in the alpine chamois. Molecular Ecology 17, 4053–4067.

Moser M, Murphy KM (2000) Dendritic cell regulation of TH1-TH2 development. Nature Immunology 1, 199–205.

Muirhead CA (2001) Consequences of population structure on genes under balancing selection. Evolution 55, 1532–1541.

Oladiran A, Belosevic M (2012) Immune evasion strategies of trypanosomes: a review. The Journal of Parasitology 98, 284–292.

Oliver MK, Telfer S, Piertney SB (2009) Major histocompatibility complex (MHC) heterozygote superiority to natural multi-parasite infections in the water vole *(Arvicola terrestris*). Proceedings of the Royal Society of London B: Biological Sciences 276, 1119–1128.

Paterson S, Wilson K, Pemberton JM (1998) Major histocompatibility complex variation associated with juvenile survival and parasite resistance in a large unmanaged ungulate population *(Ovis aries* L.). Proceeding of the National Academy of Sciences 95, 3714–3719.

Pavey SA, Sevellec M, Adam W, et al. (2013) Nonparallelism in MHCIIβ diversity accompanies nonparallelism in pathogen infection of lake whitefish *(Coregonus clupeaformis)* species pairs as revealed by next-generation sequencing. Molecular Ecology 22, 3833–3849.

Piertney SB, Oliver MK (2006) The evolutionary ecology of the major histocompatibility complex. Heredity 96, 7–21.

Rauch G, Kalbe M, Reusch TBH (2006) Relative importance of MHC and genetic background for parasite load in a field experiment. Evolutionary Ecology Research 8, 373–386.

Reusch TB, Schaschl H, Wegner KM (2004) Recent duplication and inter-locus gene conversion in major histocompatibility class II genes in a teleost, the three-spined stickleback. Immunogenetics 56, 427–437.

Reusch TBH, Langefors Å (2005) Inter- and intralocus recombination drive MHC class IIβ gene diversification in a teleost, the three-spined stickleback *Gasterosteus aculeatus*. Journal of Molecular Evolution 61, 531–541.

Reusch TBH, Wegner KM, Kalbe M (2001) Rapid genetic divergence in postglacial populations of threespine stickleback *(Gasterosteus aculeatus*): The role of habitat type, drainage and geographical proximity. Molecular Ecology 10, 2435–2445.

Revelle W (2016) psych: Procedures for Personality and Psychological Research. Northwestern University, Evanston, Illinois, USA,. http://CRAN.R-project.org/package=psych

Roche PA, Furuta K (2015) The ins and outs of MHC class II-mediated antigen processing and presentation. Nature Reviews Immunology 15, 203–2016.

Salgame P, Yap GS, Gause WC (2013) Effect of helminth-induced immunity on infections with microbial pathogens. Nat Immunol 14, 1118–1126.

Sato A, Figueroa F, O’hUigin C, Steck N, Klein J (1998) Cloning of major histocompatibility complex (Mhc) genes from threespine stickleback, *Gasterosteus aculeatus*. Molecular Marine Biology and Biotechnology 7, 221–231.

Savage AE, Zamudio KR (2011) MHC genotypes associate with resistance to a frog-killing fungus. Proceedings Of The National Academy Of Sciences Of The United States Of America 108, 16705–16710.

Schierup MH, Vekemans X, Charlesworth D (2000) The effect of subdivision on variation at multi-allelic loci under balancing selection. Genetics Research 76, 51–62.

Schwensow N, Fietz J, Dausmann KH, Sommer S (2007) Neutral versus adaptive genetic variation in parasite resistance: importance of major histocompatibility complex supertypes in a free-ranging primate. Heredity 99, 265–277.

Sebastian A, Herdegen M, Migalska M, Radwan J (2016) AMPLISAS: a web server for multilocus genotyping using next-generation amplicon sequencing data. Molecular Ecology Resources 16, 498–510.

Siddle HV, Marzec J, Cheng Y, Jones M, Belov K (2010) MHC gene copy number variation in Tasmanian devils: implications for the spread of a contagious cancer. Proceedings Of The Royal Society B-Biological Sciences 277, 2001–2006.

Sinigaglia F, Guttinger M, Kilgus J, et al. (1988) A malaria T-cell epitope recognized in association with most mouse and human MHC class II molecules. Nature 336, 778–780.

Slade RW, McCallum HI (1992) Overdominant vs. frequency-dependent selection at MHC loci. Genetics 132, 861–864.

Spurgin LG, Richardson DS (2010) How pathogens drive genetic diversity: MHC, mechanisms and misunderstandings. Proceedings of the Royal Society B: Biological Sciences 277, 979–988.

Stuart YE, Veen T, Weber JN, et al. (In review) Environmental explanation for semi-parallel evolution. Nature Ecology and Evolution.

Stutz WE, Bolnick DI (2014) A Stepwise Threshold Clustering (STC) method to infer genotypes from error-prone next-generation sequencing of multi-allele genes such as the Major Histocompatibility Complex (MHC). PLoS ONE.

Sutton JT, Nakagawa S, Robertson BC, Jamieson IG (2011) Disentangling the roles of natural selection and genetic drift in shaping variation at MHC immunity genes. Molecular Ecology 20, 4408–4420.

Takahata N, Nei M (1990) Allelic genealogy under overdominant and frequency-dependent selection and polymorphism of major histocompatibility complex loci. Genetics 124, 967–978.

Takahata N, Satta Y, Klein J (1992) Polymorphism and balancing selection at Major Histocompatibility Complex loci. Genetics 130, 925–938.

Thursz MR, Kwiatkowski D, Allsopp CE, et al. (1995) Association between an MHC class II allele and clearance of hepatitis B virus in the Gambia. New England Journal of Medicine 332, 1065–1069.

Tobler M, Plath M, Riesch R, et al. (2014) Selection from parasites favours immunogenetic diversity but not divergence among locally adapted host populations. Journal of Evolutionary Biology 27, 960–974.

Warrens MJ (2008) On association coefficients for 2×2 tables and properties that do not depend on the marginal distributions. Psychometrika 73, 777–789.

Weber J, Bradburd GS, Stuart YE, Stutz WE, Bolnick DI (2017) The relative contributions of distance, landscape resistance, and habitat, to genomic divergence between parapatric lake and stream stickleback. Evolution **Online Early**.

Wedekind C, Walker M, Little TJ (2006) The separate and combined effects of MHC genotype, parasite clone, and host gender on the course of malaria in mice. BMC Genetics 7, 55.

Wegner K, Kalbe M, Schaschl H, Reusch T (2004) Parasites and individual major histocompatibility complex diversity?an optimal choice? Microbes and Infection 6, 1110–1116.

Wegner KM, Kalbe M, Kurtz J, Reusch TBH, Milinski M (2003) Parasite selection for immunogenetic optimality. Science 301, 1343.

Wegner KM, Kalbe M, Milinski M, Reusch TB (2008) Mortality selection during the 2003 European heat wave in three-spined sticklebacks: effects of parasites and MHC genotype. BMC Evolutionary Biology 8, 124.

Westerdahl H, Asghar M, Hasselquist D, Bensch S (2012) Quantitative disease resistance: to better understand parasite-mediated selection on major histocompatibility complex. Proceedings of the Royal Society of London B: Biological Sciences 279, 577–584.

Yasukochi Y, Satta Y (2013) Current perspectives on the intensity of natural selection of MHC loci. Immunogenetics 65, 479–483.

